# Synergistic *MAPT* mutations as a platform to uncover modifiers of tau pathogenesis

**DOI:** 10.1101/2025.02.07.636933

**Authors:** Miles R. Bryan, Michael F. Almeida, Kyle Pellegrino, Carli K. Opland, J. Ethan Paulakonis, Jake McGillion-Moore, Hanna Trzeciakiewicz, Diamond King, Xu Tian, Jui-Heng Tseng, Jonathan C. Schisler, Nicholas G. Brown, Ben A. Bahr, Todd J. Cohen

## Abstract

The natively unfolded tau (MAPT) protein is extremely soluble, which poses challenges when modeling neurofibrillary tangle (NFT) pathology in Alzheimer’s disease (AD). To overcome this hurdle, we combined P301L and S320F mutations (PL-SF) to generate a rapid and reliable tau pathology platform to expedite the discovery of factors that modify tau aggregation. Using this model, we evaluated heat-shock proteins (Hsp), which have been linked to tau pathology, but whose role in AD remains enigmatic and controversial. In primary neurons, expression of Hsp70, but not Hsc70 or Hsp90, exacerbated tau aggregation. Conversely, lowering Hsp70 or employing a chaperone-deficient tau mutant (PL-SF-4Δ) reduced tau phosphorylation and abrogated tau aggregation, highlighting Hsp70 as a key driver of tau aggregation. Hsp70 foci clustered within and surrounding neuritic plaques and NFTs in post-mortem AD brain. Functionally, mature aggregate-bearing neurons showed deficits in neuronal firing and network communication, which were restored by chaperone-binding deficient tau variants that abrogated tau pathology. This study provides a powerful cell-intrinsic model for accelerated tau aggregation, which can be harnessed to identify potent regulators of tau aggregation as promising therapeutic targets.

## Introduction

Aggregation of the microtubule-associated protein tau (MAPT) and the formation of insoluble tau inclusion pathology are hallmark features of Alzheimer’s disease and related neurodegenerative disorders. In contrast to AD, in which tau pathology is comprised of normal wild-type tau proteins, pathogenic disease-causing mutations in *MAPT* are implicated in familial frontotemporal dementia linked to chromosome 17 (FTDP-17). Over 50 different pathogenic *MAPT* missense, silent, and intronic mutations have been identified^1^. In many instances, missense mutations generate loss and gain of function phenotypes including altered microtubule (MT) binding and enhanced tau aggregation propensity, which are implicated in prion-like tau spreading along defined spatial and temporal brain networks^2–4^.

While familial missense mutations at residue P301 (e.g., P301L and P301S) have been well characterized as aggregate-prone and MT-binding deficient, less is known about nearby mutations within the MT-binding repeat domain (MTBR)^5,6^. Our recent study characterized a panel of FTDP-17 tau mutations spanning this domain and found that the cellular surveillance machinery composed of HDAC6 and associated chaperones (e.g. Hsp70) shows increased binding to two specific mutations within the R3 domain (P301L and S320F), but not others, suggesting these two mutants are conformationally distinct and are more effectively recognized by the quality control machinery that targets tau for degradation^7^.

In separate studies, the P301L and S320F mutations were combined into the same full-length tau molecule (termed PL-SF), which generated a surprisingly aggregate-prone conformation leading to an accelerated AD-like tangle pathology, neuroinflammation, and memory impairments in mice and rhesus macaques^8–12^. Thus, PL-SF could potentially be leveraged as an intrinsic, cell-autonomous, and seeding-independent model of full-length tau aggregation, which would otherwise be challenging to achieve with full-length tau. In fact, many widely used tau models, including the tau biosensor cell lines, employ artificially truncated tau mutants comprising only the tau repeat domain, a fragment with questionable physiological relevance to neurons^13^. In comparison, the PL-SF model offers several advantages, including serving as a rapid model of full-length and mature tau aggregation as a cornerstone for high-throughput discovery of tau modifiers^8,9,11,14^.

Among known tau interactors, heat shock proteins (Hsps) such as inducible Hsp70, constitutive Hsc70and related Hsp90 have high affinity for aberrant forms of tau, yet their functions as refolding and/or triaging factors remains unclear^8,9,15,16–25^. This debate has obvious therapeutic implications since Hsp inhibitors are in clinical development and have been evaluated in preclinical models of AD^26–31^. However, prior studies have yielded quite contradictory and even opposing results, since on the one hand, Hsp70 is thought to promote the MC1-positive tau conformation^18^, while other studies suggest Hsp70 and Hsc70 mediate tau degradation via autophagy and the ubiquitin-proteasome system (UPS)^32,33^. One possible explanation for these discrepancies is the inability to generate mature tau pathology in these studies, as observed in human AD brains. Adding additional complexity, tau interacts with dozens of chaperone adaptors and cofactors that may promote or suppress chaperone function directed towards tau^20,21,34,35^. Here, we characterized a tau PL-SF model of accelerated tau pathology, which allowed us to interrogate the role of Hsp70 and other chaperones as bona fide modifiers of tau-mediated toxicity and aggregation.

## Results

### Synergistic familial *MAPT* mutations drive tau phosphorylation and aggregation

We first evaluated tau aggregation in PL-SF expressing immortalized QBI-293 cells. Cells transfected with human T40 (2N4R) wild-type, P301L, S320F, or PL-SF were monitored for RIPA-soluble expression of phosphorylated tau (AT8 and AT100) and cleaved tau (TauC3), two well-established markers of tau pathogenesis. Phosphorylated tau accumulated in PL-SF expressing cells, more so than WT or either single mutant (**Figure 1 A, B**). Confirming this finding, total tau immunoblotting revealed up-shifted tau protein bands in SF and PL-SF, indicative of tau hyperphosphorylation. Surprisingly, the single PL mutant appeared hypo-phosphorylated compared to WT or SF mutations. Immunocytochemistry confirmed these findings and revealed Thioflavin S (ThioS) positive SF and PL-SF aggregates, consistent with enhanced tau aggregation (**Supplemental Figure 1, Figure 1C**). Similar to phosphorylated tau, cleaved tau also preferentially accumulated in SF or PL-SF expressing cells.

**Figure 1:**
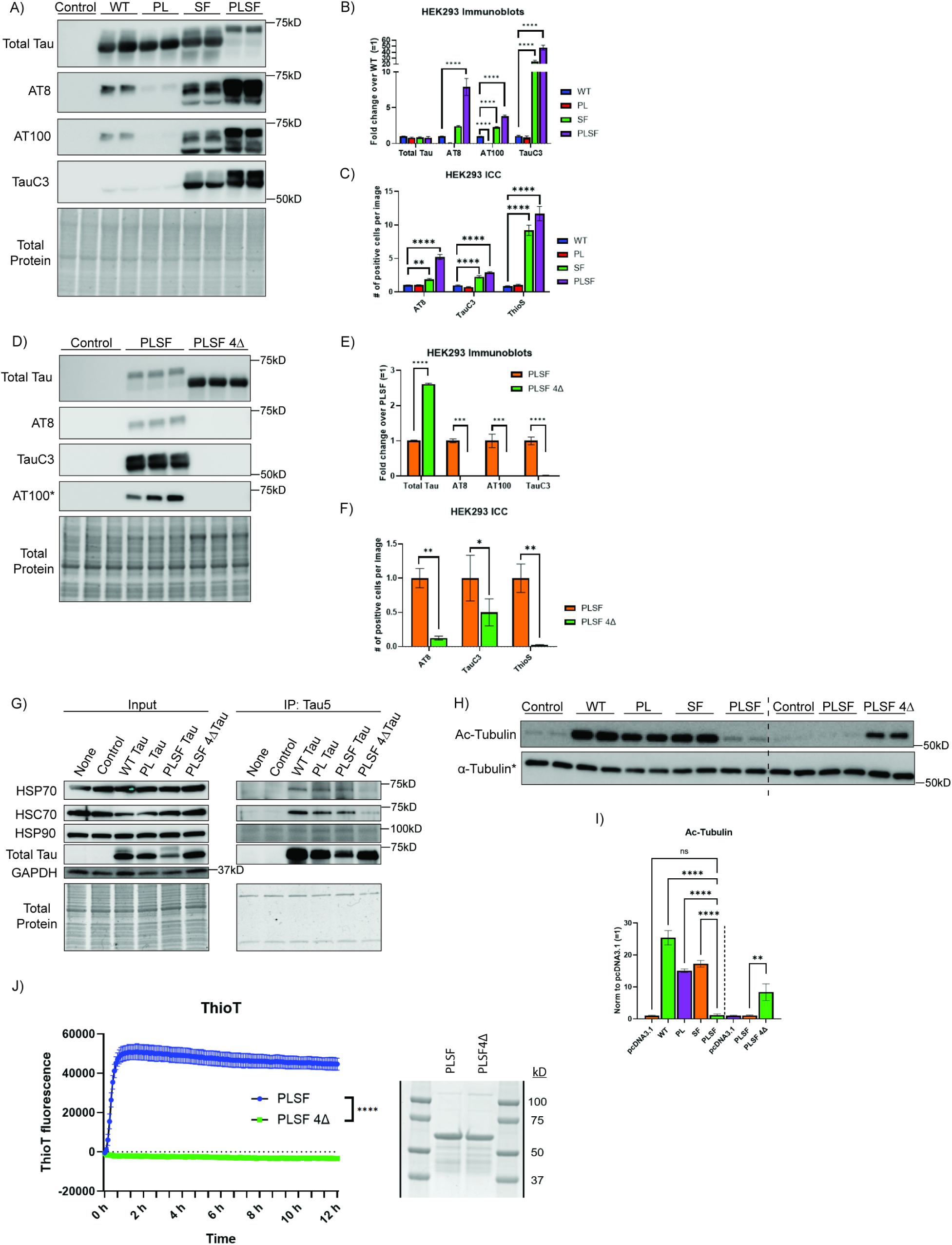
Abrogating tau binding to heat shock proteins alleviates pathology in a PL-SF tau aggregation model. **A)** Representative immunoblots (RIPA fraction) of QBI-293 cells transfected with 2N4R WT, P301L, S320F, or P301L-S320F tau. **B)** Immunoblot quantification of total tau, p-tau (AT8 and AT100), and cleaved tau (TauC3) in RIPA fraction. N=4 per condition; **C)** Immunocytochemistry quantification of QBI-293 cells transduced with WT, P301L, S320F, or PL-SF tau labeled with AT8, TauC3, or ThioS. N=48 FOVs per condition. *= p-value significant by one-way ANOVA. Representative images found in Supplemental Figure 1. **D)** Representative immunoblots (RIPA fraction) of QBI-293 cells transfected with 2N4R P301L-S320F tau or P301L-S320F 4Δ (deletion of I277/278, I308/V309 residues). *AT100 was run on a separate blot with the exact same lysates to improve detection. **E)** Immunoblot quantification of total tau, p-tau (AT8 and AT100), and cleaved tau (TauC3) in RIPA soluble fractions. N=3 per condition. **F)** Immunocytochemistry quantification of QBI-293 cells transduced with 2N4R PL-SF or PL-SF 4Δ tau labeled with AT8, TauC3, or ThioS. N=16 FOVs per condition. *= p-value significant by unpaired Student’s t-test. Representative images found in Supplemental Figure 3. **G)** Co-immunoprecipitation of QBI-293 cells transfected with WT, P301L, PL-SF, or PL-SF 4Δ tau. Tau5 antibody was used to IP tau and immunoblots were probed with total tau (rabbit polyclonal) and Hsp70, Hsc70, and Hsp90 antibodies. **H)** Representative immunoblots of QBI-293 cells transfected with 2N4R tau variants and probed with acetyl-tubulin and total alpha-tubulin antibodies. Note: Acetyl tubulin and alpha tubulin blots were run separately with the same lysates to improve detection. **I)** Quantification of acetyl-tubulin across all 2N4R tau variants. -- line signifies separate sets of transfections in panel I and J. N=4 per condition; *= p-value significant by one-way ANOVA. Control= empty plasmid vector (pcDNA3.1)**. J)** Quantification of in vitro Thioflavin T aggregation assay using purified protein from PLSF and PLSF 4Δ. N=4 replicates per condition. *= significant by unpaired student t-test at 12hr timepoint. Coomassie (right) shows equal protein loading into reactions.

Multiple familial FTD-associated mutations occur at residue P301 (P301L, P301S, P301T) and S320 (S320F, S320Y). We evaluated a panel of other combination mutants by immunoblotting and immunocytochemistry and found that all double mutants formed phosphorylated, cleaved, and insoluble tau species compared to individual single tau mutants (**Supplemental Figure 2**). The most consistently insoluble tau species was PL-SF, which we selected for follow-up analysis.

A prior study found that deletion of four key residues in tau (I277/I278 and I308/V309) can abrogate Hsp70 binding^36^. Since the role of Hsp70 in tau pathology is unclear, we generated a PL-SF mutant in which these four residues were depleted, termed PL-SF 4Δ. In transfected QBI-293 cells, PL-SF 4Δ was nearly completely dephosphorylated and accumulated negligible amounts of cleaved tau. PL-SF 4Δ was also downshifted compared to PL-SF (**Figure 1D, E**) and showed significantly less phosphorylated (AT8), cleaved (TauC3), and aggregated (ThioS) tau (**Supplemental Figure 3, Figure 1F**). By co-immunoprecipitation assays, we confirmed that PL-SF 4Δ abrogated the majority of Hsc70 and Hsp70 binding to tau. We did not observe detectable and specific binding to endogenous Hsp90 in our model (**Figure 1G**).

We also assessed the impact of tau mutants on acetylated-tubulin (ac-tubulin), a readout for microtubule (MT) stability and dynamics. Wild-type tau increased ac-tubulin, as expected, while P301L or S320F were less effective (∼30-40%). Consistent with a near-complete loss of MT regulatory function, PL-SF reduced ac-tubulin levels down to the levels of non-transfected cells. Surprisingly, PL-SF 4Δ partly restored the tubulin interactions (**Figure 1H, I**).

To confirm that PL-SF has increased intrinsic propensity to self-aggregate, as opposed to being influenced by other cellular factors, we performed in vitro Thioflavin T (ThioT) assays using recombinant purified tau proteins to assess aggregation kinetics in the presence of heparin as an inducing agent. Indeed, PL-SF tau rapidly aggregated within 1 hr post-induction, at which point the formation of recombinant tau fibrils plateaued, while PL-SF 4Δ was completely resistant to aggregation, even though similar tau concentrations were present (**Figure 1J**).

### Impaired chaperone binding alleviates tau pathology in neurons

To validate our findings in neurons, we developed mutant tau lentiviral constructs suitable for high-efficiency transduction into mouse primary cortical neurons. Transduced primary neurons were evaluated with a panel of tau mutants, including PL-SF. Immunoblotting of neuronal lysates showed that PL-SF tau was again up-shifted and accumulated more phosphorylated tau (**Figure 2A, B).** Unlike immortalized cell lines, neurons did not show appreciable levels of cleaved tau (TauC3). Only PL-SF accumulated in the insoluble fraction (**Figure 2C**). PL-SF 4Δ showed a marked reduction in tau phosphorylation and aggregation, suggesting a similar phenotype in neurons as well. In fact, deletion of just two of the Hsp-binding residues (ΔI277/I278 or ΔI308/V309) abrogated tau phosphorylation and aggregation, with residues I308/V309 showing the most dominant effect (**Figure 2A-C**). Introduction of β-sheet disrupting proline mutations has been shown to abrogate tau aggregation^37–41^. Indeed, by mutating residues I277 and I308 to prolines (PL-SF I277P/I308P), we were able to suppress tau aggregation to similar levels as seen in PL-SF 4Δ (**Supplemental Figure 4**).

**Figure 2:**
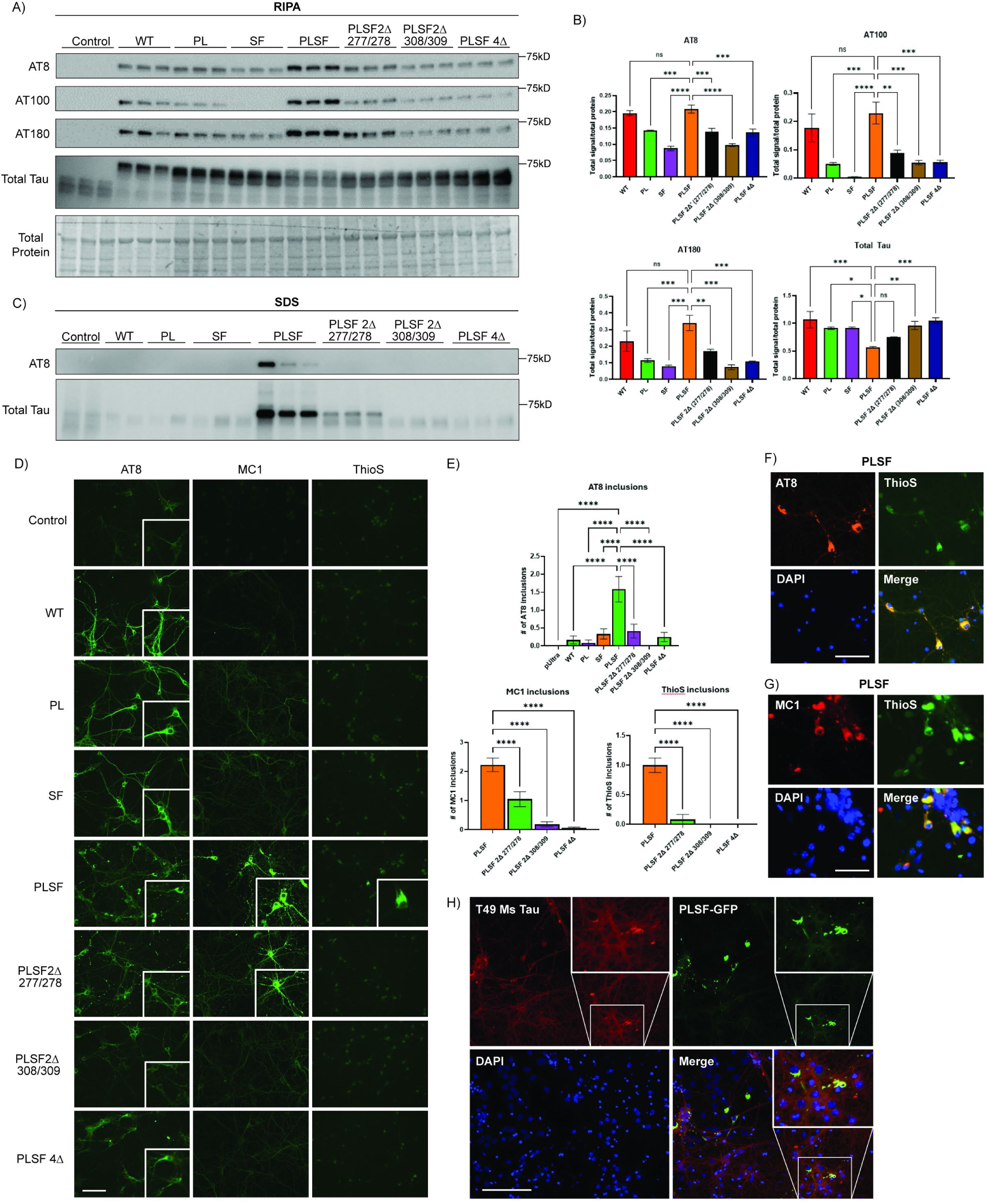
Development and characterization of a PL-SF tau aggregation model in primary cortical neurons. **A)** Representative RIPA-soluble immunoblots of DIV10 primary cortical neurons transduced with lentiviruses expressing 2N4R WT, P301L, S320F, P301L-S320F, or PL-SF 2Δs (R2 and R3) and 4Δ tau variants. **B)** Quantification of AT8, AT100, AT180, and total tau immunoblots in primary cortical neurons (RIPA fraction). N=3 per condition; *= p-value significant by one-way ANOVA. **C)** Representative SDS-soluble immunoblots of DIV10 primary cortical neurons transduced with lentiviruses expressing 2N4R WT, P301L, S320F, P301L-S320F, or PL-SF 2Δ and 4Δ tau variants. N=2 for control, WT, and P301L; N=3 for other variants. **D)** Representative immunocytochemistry images of 2N4R tau variants labeled with AT8 (p-tau), MC1 (conformational tau), ThioS (aggregated tau) in GFP. Scale bar= 125 µm. **E)** Quantification of the number of AT8, MC1, and ThioS inclusions in PL-SF and PL-SF 2Δ (R2 and R3) and 4Δ variants. N=12-24 FOVs per condition. *= significant by one-way ANOVA. **F, G)** Representative immunocytochemistry images of PL-SF tau displaying colocalization between F) AT8 (p-tau) or G) MC1 (conformational tau) with ThioS (aggregated tau). Scale bar= 50um. **H)** Representative immunocytochemistry of PL-SF GFP colocalized with T49 tau (mouse-specific). Scale bar= 125 µm. Control= empty viral vector (pUltra).

By performing microscopy on transduced neurons, we found that PL-SF accumulated AT8, MC1, and ThioS-positive tau inclusions (**Figure 2D, E**). Both MC1 (pre-tangle confirmational tau) and AT8 immunoreactive tau were amorphous and globular and accumulated as large varicosities, or swellings, within neurites. The large globular tau aggregates colocalized with ThioS, supporting tangle-like pathology (**Figure 2F, G**). By transducing neuronal cultures with GFP-tagged PL-SF and labeling neurons with a mouse-specific tau antibody (monoclonal T49), we observed recruitment of endogenous mouse tau to PL-SF inclusions, implying that no species barrier exists between human PL-SF aggregates and mouse tau (**Figure 2H**). PL-SF behaved similarly in tau knockout neurons, and therefore, endogenous tau is not required for the accumulation of PL-SF pathology in neurons (**Supplemental Figure 5**).

### Hsp70 promotes tau hyperphosphorylation and aggregation

To determine whether Hsp70 is a more potent regulator of tau pathology compared to other tau-associated heat shock proteins, PL-SF expressing neurons were co-transduced with either HSP70, HSC70, or HSP90. Only Hsp70, but not Hsc70 or Hsp90, exacerbated tau phosphorylation (AT8), which was supported by an appreciable upshift in tau’s molecular weight (**Figure 3A, B**) and a trend towards increased levels of total insoluble tau (**Figure 3C, D**). Hsp70 significantly increased AT8, MC1, and ThioS immunoreactivity (**Figure 3E, F**). The effect of Hsp70 appears remarkably specific to the aggregate-prone PL-SF tau, since the P301L single mutant was unaffected **(Supplemental Figure 6).**

**Figure 3:**
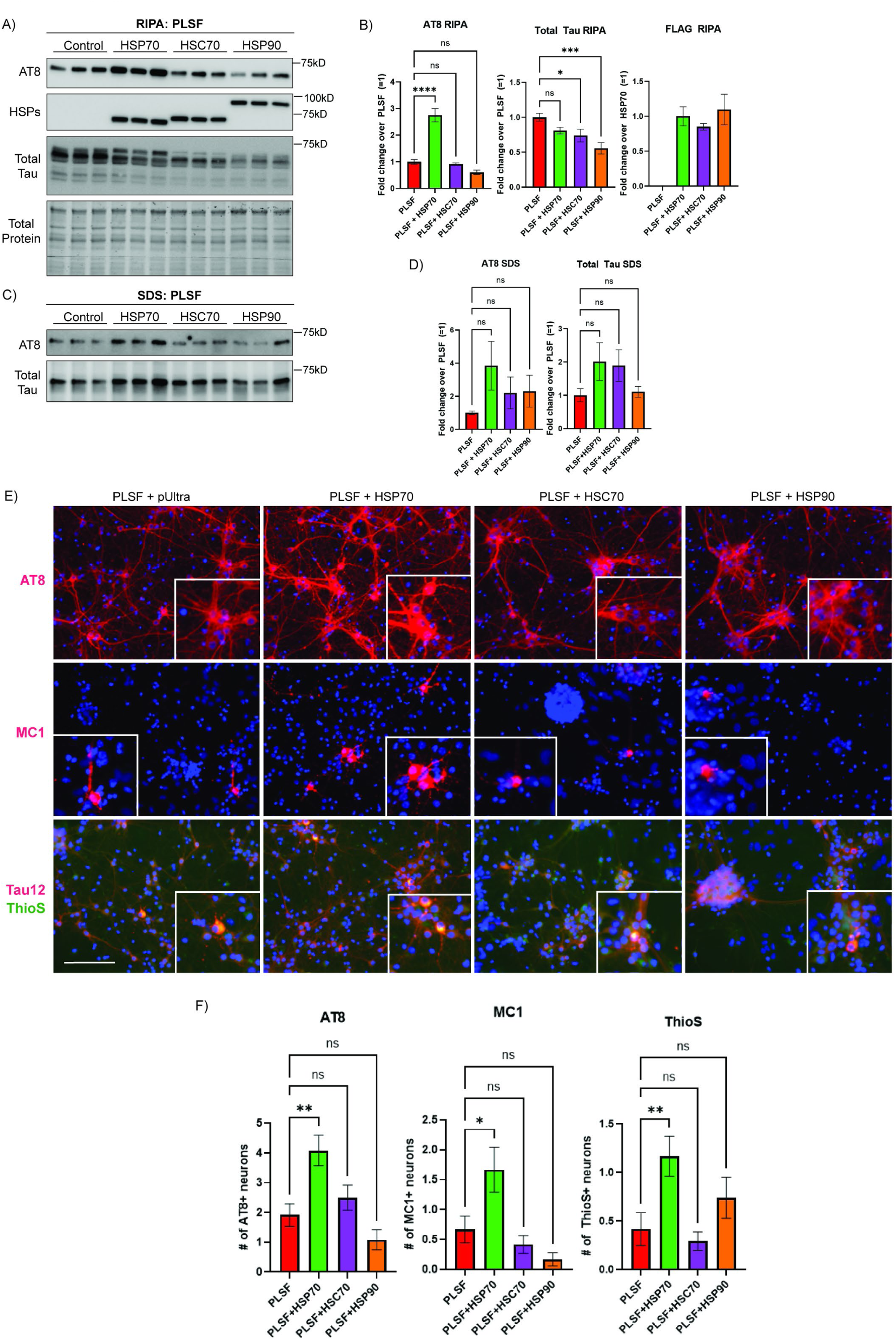
Hsp70 exacerbates tau hyperphosphorylation and aggregation. **A)** Representative immunoblots of 2N4R PL-SF tau co-transduced with HSP70, HSC70, and HSP90 in DIV10 primary cortical neurons (RIPA fraction). **B)** Quantification of AT8 (p-tau), total tau (rabbit polyclonal), or Hsp70, Hsc70, Hsp90 (FLAG-tagged) in the RIPA fraction. N=6 per condition. *= p-value significant by one-way ANOVA. **C)** Representative immunoblots of 2N4R PL-SF tau co-transduced with HSP70, HSC70, and HSP90 in DIV10 primary cortical neurons (SDS fraction). **D)** Quantification of AT8 (p-tau), total tau (rabbit polyclonal) in the SDS fraction. N=6 per condition. ns= p-value is not significant by one-way ANOVA. **E)** Representative immunocytochemistry images of primary cortical neurons co-transduced 2N4R PL-SF tau and HSP70, HSC70, or HSP90 labeled with AT8, MC1, Tau12 (RFP) or ThioS (GFP). Scale bar= 125 µm. **F)** Quantification of MC1, AT8, and ThioS positive inclusions in primary cortical neurons co-transduced 2N4R PL-SF tau and either HSP70, HSC70, or HSP90. n=12 FOVs per condition. *= p-value significant by one-way ANOVA. Control= empty viral vector (pUltra).

Next, we depleted HSP70 in PL-SF transduced neurons. We achieved ∼ 50% Hsp70 knockdown using lentiviral shRNA transduction, which did not affect HSC70 expression. Strikingly, HSP70 knockdown led to ∼70% decrease in phosphorylated tau (**Figure 4 A, B**) and ∼50% decrease in total insoluble tau (**Figure 4C, D**). We note that a ∼50% decrease in soluble tau was also observed, suggesting that Hsp70 depletion enhances the turnover of steady-state tau levels (**Figure 4A, B**). The impact of Hsp70 on tau also appears to be quite specific since knockdown of HSP90 did not alter total or phosphorylated tau (**Supplementary Figure 7**). Overall, our findings highlight Hsp70 as a primary regulator of aggregate-prone tau species in this accelerated PL-SF tau model.

**Figure 4:**
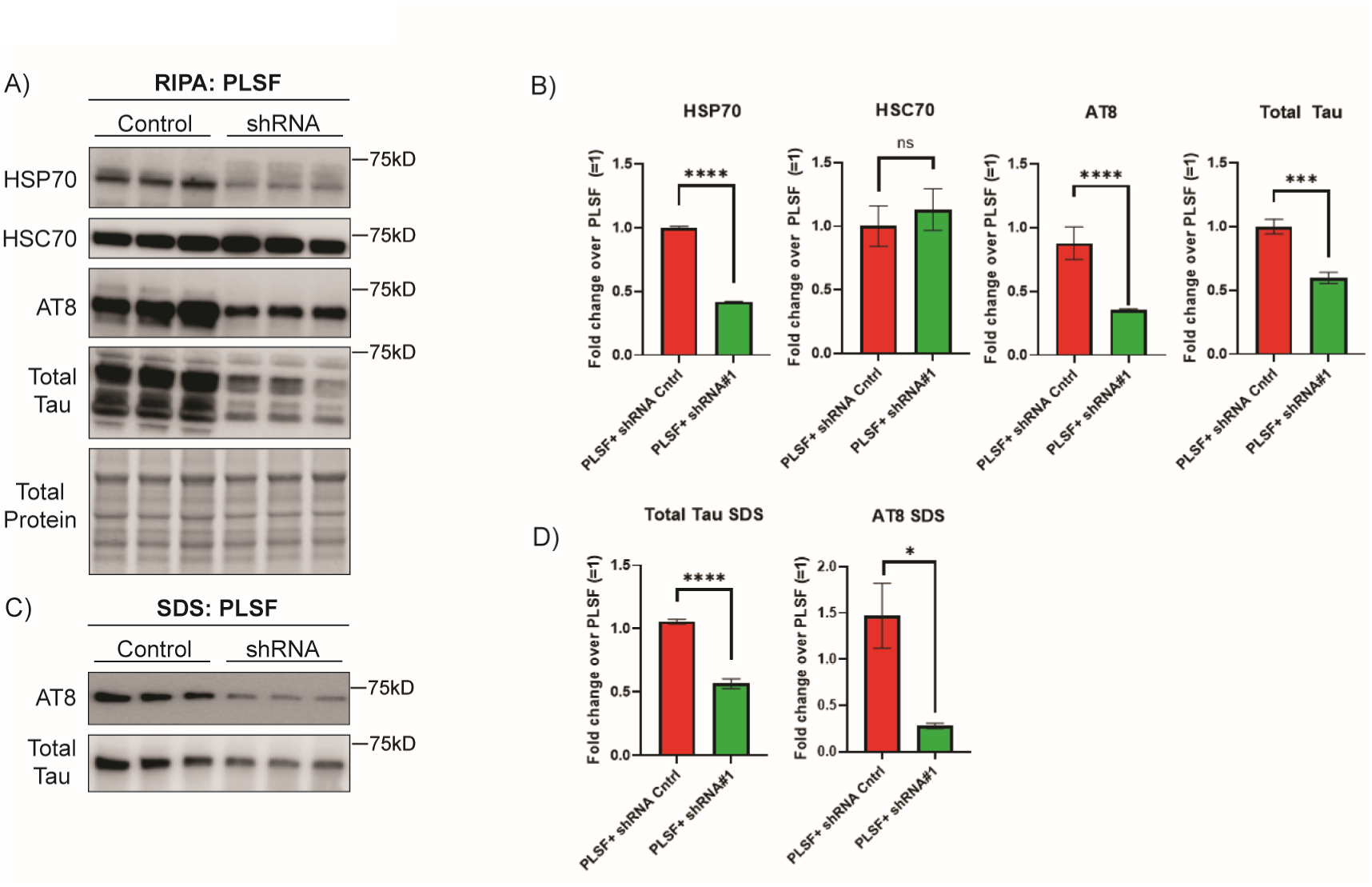
Reducing Hsp70 can suppress tau hyper-phosphorylation and aggregation. **A)** Representative immunoblots of 2N4R PL-SF tau co-transduced with a shRNA against HSP70 in DIV10 primary cortical neurons (RIPA fraction). **B)** Quantification of AT8 (p-tau), total tau (rabbit polyclonal) and endogenous Hsp70 and Hsc70 in the RIPA fraction. N=3. **C)** Representative immunoblots of 2N4R PL-SF tau co-transduced a shRNA against HSP70 in DIV10 primary cortical neurons (SDS fraction). **D)** Quantification of AT8 (p-tau), total tau (rabbit polyclonal) in the SDS fraction. N=3. *= p-value significant by unpaired Student’s t-test. Control= empty viral vector (pUltra).

### Cell-autonomous aggregated tau alters neuronal activity and network synchrony

Given the extensive tau pathology observed in PL-SF transduced neurons, we evaluated any deficits in neuronal activity and connectivity. Electrically active neurons were transduced at DIV18 following an initial multi-electrode array (MEA) recording. Daily recordings were taken for 53 days, and in parallel, neurons were immunolabeled at DIV29 and DIV62 to assess the extent of tau pathology.

We evaluated the mean firing rate (a measure of overall activity), the number of network bursts (a measure of neuronal connectivity), and the area under the normalized crossed-correlation (a readout for neuronal synchrony). After 15-20 days post-lentiviral transduction, PL-SF neurons showed a decline in all three metrics, suggesting a loss in neuronal activity and reduced synchronous firing across neuronal networks. The deficits worsened over time, but some neurons continued to fire even past DIV50 (**Figure 5A-C**). Raster plots at DIV54 highlight reduced network firing across electrodes and elongated network firing time (**Figure 5D-F**). PL-SF 4Δ did not exhibit any deficits in any of the activity metrics and showed similar firing patterns as compared to wild-type control neurons (**Figure 5A-C**).

**Figure 5:**
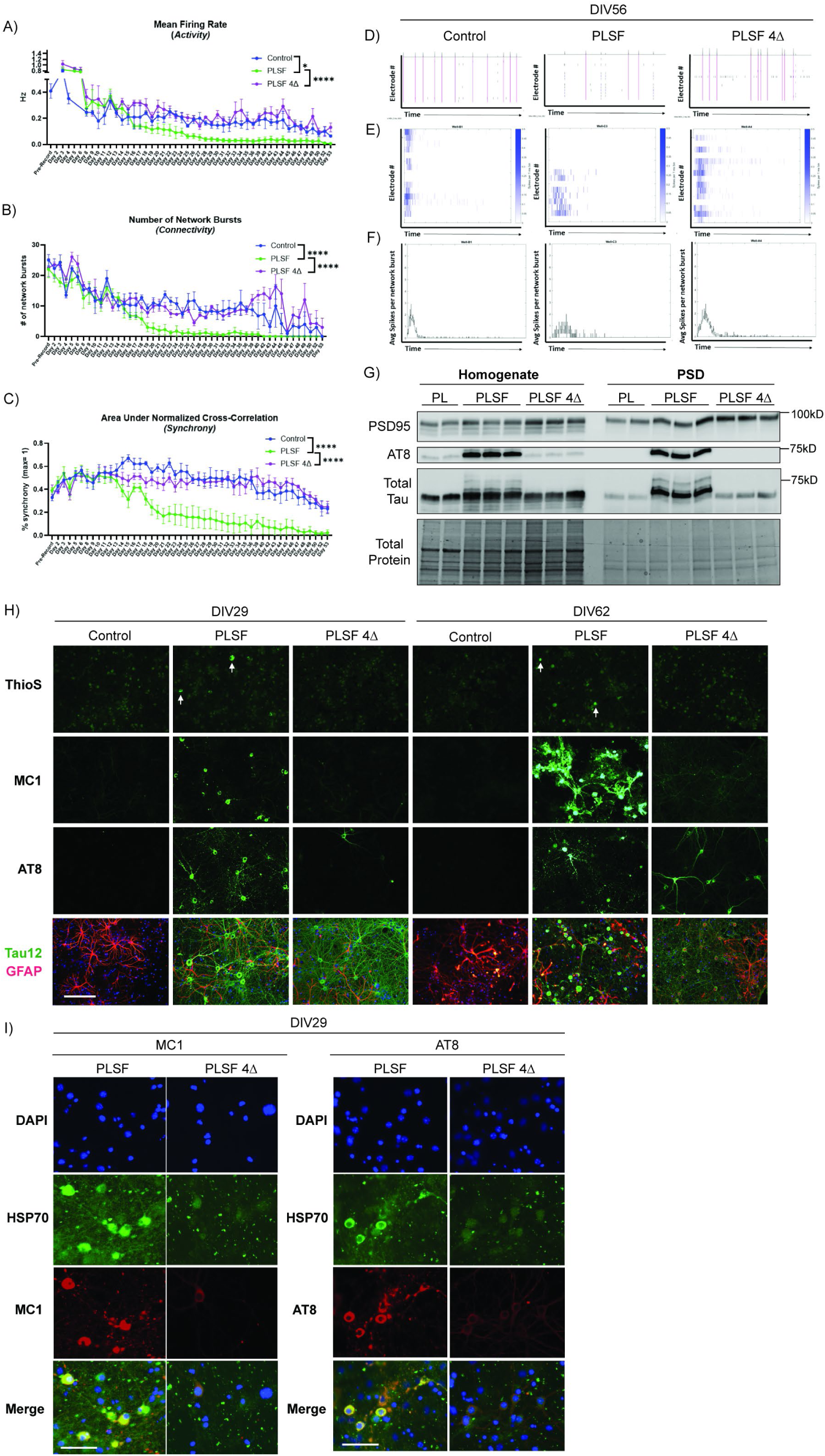
Mature tau aggregates cause functional deficits in neurons that are reversed by abrogating tau-Hsp binding. **A-C)** Quantification of neuronal activity comparing PL-SF to PL-SF 4Δ across 53 days of multi-electrode array recording starting at DIV18 following initial pre-recording. A) Mean firing rate-activity B) Number of network bursts-connectivity C) Area under normalized cross-correlation-synchrony. N=4 replicate wells per condition. *= p-value significant by one-way ANOVA. **D-F)** Representative raster and network burst plots at DIV54 (day 36 of recording) comparing empty control (pUltra) to PL-SF and PL-SF 4Δ. D) Number of network bursts across electrodes over time (pink lines) E) number of spikes across electrodes over time F) Average number of spikes per network burst over time. **G)** Representative immunoblot of isolated post synaptic density (PSD) in DIV18 primary cortical neurons transduced with lentiviruses expressing P301L, PL-SF, or PL-SF 4Δ. Immunoblot was probed with PSD95, AT8 (p-tau), and rabbit polyclonal tau (total tau). **H)** Representative 20X immunocytochemistry images of primary cortical neurons comparing empty vector control (pUltra) to PL-SF and PL-SF 4Δ tau variants at DIV29 and DIV62. Neurons were labeled with ThioS (aggregated tau), MC1 (conformational tau), AT8 (p-tau), and Tau12 (total tau) in GFP. Tau12 was co-labeled with GFAP (RFP) to label astrocytes. Scale bar= 125 µm. **I)** Representative immunocytochemistry images of primary cortical neurons comparing PL-SF and PL-SF 4Δ at DIV29. Neurons were labeled with endogenous Hsp70 (GFP), MC1 or AT8 (RFP), and DAPI (BFP). Scale bar= 50 µm. Control= empty viral vector (pUltra).

The impact of aggregate-prone tau mutants on neuronal firing could represent a direct effect of tau at the synapse. Therefore, we assessed whether PL-SF accumulated at the post-synaptic density (PSD), a re-localization thought to coincide with synaptic dysfunction. PSD fractions were biochemically isolated from transduced neurons at DIV18 and probed by immunoblotting for phosphorylated and total tau. PL-SF accumulated in the PSD fraction, which was not observed with PL-SF 4Δ or any single tau mutants (e.g., P301L) (**Figure 5G**). We conclude that mature PL-SF tau aggregates accumulate at the PSD and drive synaptic dysfunction. By abrogating Hsp binding, PL-SF 4Δ bypasses these defects.

PL-SF expressing neurons were aged in culture to determine tau phenotypes over time. Indeed, aged neurons accumulated dense tau inclusions comprised of phosphorylated, MC1, and ThioS-positive tau at DIV29, which became even more intensely labeled by DIV62. All indicators of tau pathology were absent in PL-SF 4Δ expressing neurons, even at late time points (**Figure 5H**). We also note that Hsp70 began to colocalize with aggregated tau in aged neurons (**Figure 5I**), supporting aggregation-dependent recruitment of Hsp70.

### Cell-autonomous tau pathology in organotypic brain slice cultures

To validate our findings in brain culture systems comprised of both neurons and glia, we transduced adult rat organotypic hippocampal brain slices ex vivo with either GFP alone, wild-type tau-GFP, PL-SF-GFP, or PL-SF 4Δ-GFP at DIV1 and imaged brain slices after nine weeks in culture **(Figure 6A, B**). Similar to cultured neurons, PL-SF accumulated dense AT8 and MC1-positive inclusions and distinct neuritic puncta, or foci. Most tau pathology preferentially accumulated in the dentate gyrus (DG) and extended through the hippocampal CA1 and CA3 regions. In contrast, PL-SF 4Δ exhibited diffuse, non-aggregated neuronal labeling in the DG and showed far lower levels of pathogenic tau markers, such as MC1 and AT8 **(Figure 6C, D)**. These data suggest that, in a multicellular model with complete hippocampal architecture, abrogating tau-Hsp binding can confer neuroprotection.

**Figure 6:**
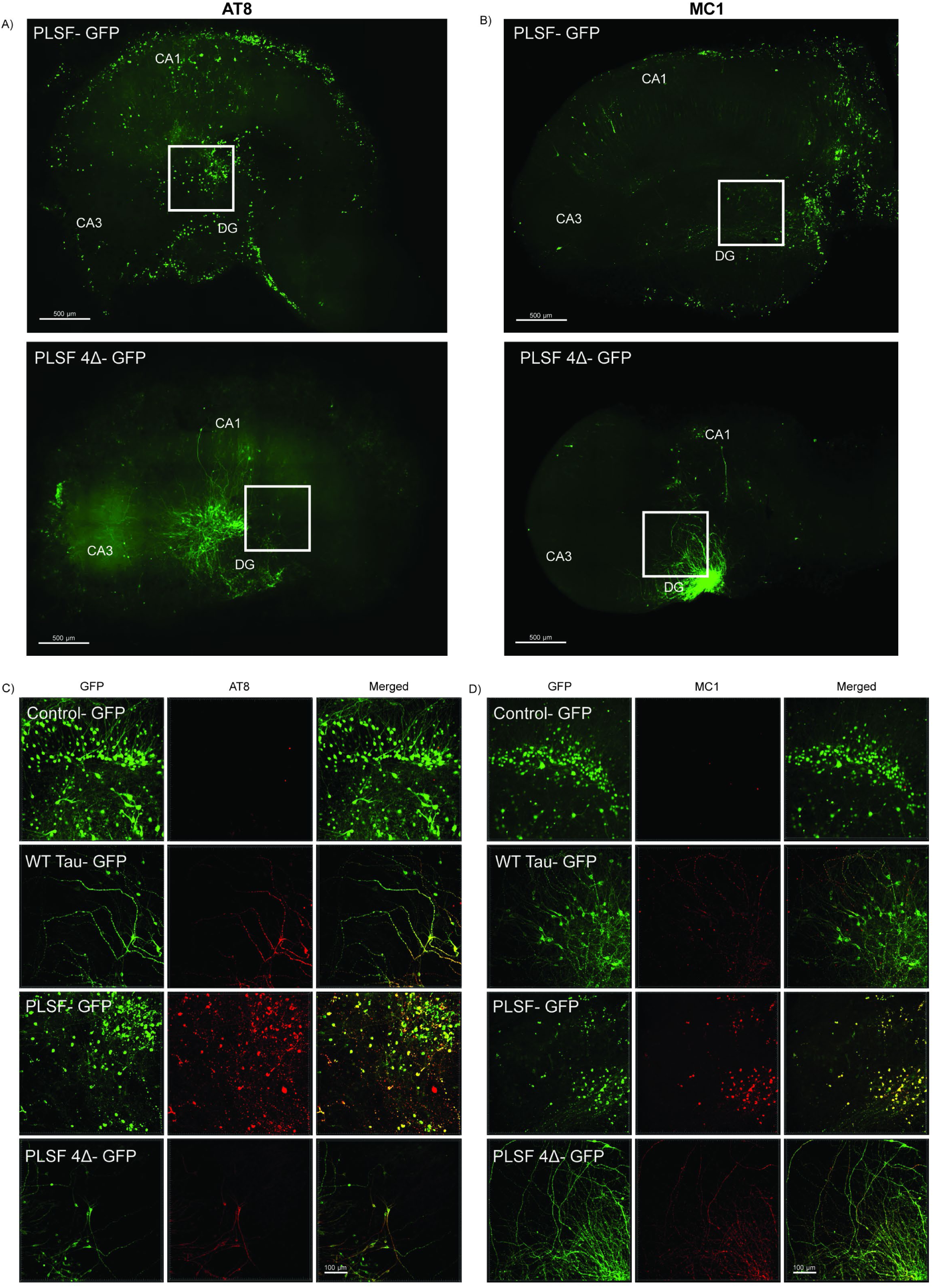
Abrogating tau-Hsp binding suppresses tau pathology in three-dimensional organotypic hippocampal slice cultures. Rat hippocampal slices were processed for immunocytochemistry after 9 weeks in culture following lentiviral infection of eGFP empty vector control, wild-type tau, PL-SF tau, and PL-SF 4Δ. **A, B)** Immunocytochemistry of the entire hippocampal slice following labeling of reveal sufficient expression of PL-SF and PL-SF 4Δ lentiviruses (GFP). Left-AT8 staining; Right-MC1 staining. CA1, CA3, and DG regions are labeled in white font. White boxes denote magnified images in panel C, D. Scale bar= 500 µm. **C, D)** Representative immunocytochemistry images of C) AT8 and D) MC1 labeled in RFP for empty vector control, wild-type tau, PL-SF tau, and PL-SF 4Δ taken from the dentate gyrus (DG) region of the hippocampal slice culture (inset white boxes from A, B). Scale bar= 100 µm.

### Hsp70 is upregulated in human AD brain and intercalates within tau aggregates

Finally, we evaluated Hsp70 levels and localization in AD human post-mortem brain. Biochemical fractionation of control and AD brain lysates showed elevated Hsp70 expression in both soluble (hi-salt extracted) and insoluble (urea extracted) whole cortex AD brain homogenates (**Figure 7A,B**). Hsc70 was only upregulated in the insoluble fraction of AD brains (**Figure 7C,D**). We then performed immunohistochemistry on AD hippocampus and found that the vast majority of AT8-positive tau aggregates (plaques and tangles) were labeled by Hsp70-positive foci (**Figure 7E, Supplemental Figure 8**). Interestingly, the Hsp70 foci formed discrete puncta that appeared to intercalate within the AT8-positive tau neuritic plaques and neurofibrillary tangles (NFTs). This data further supports a prominent role for Hsp70 in modulating tau accumulation in human AD brain.

**Figure 7:**
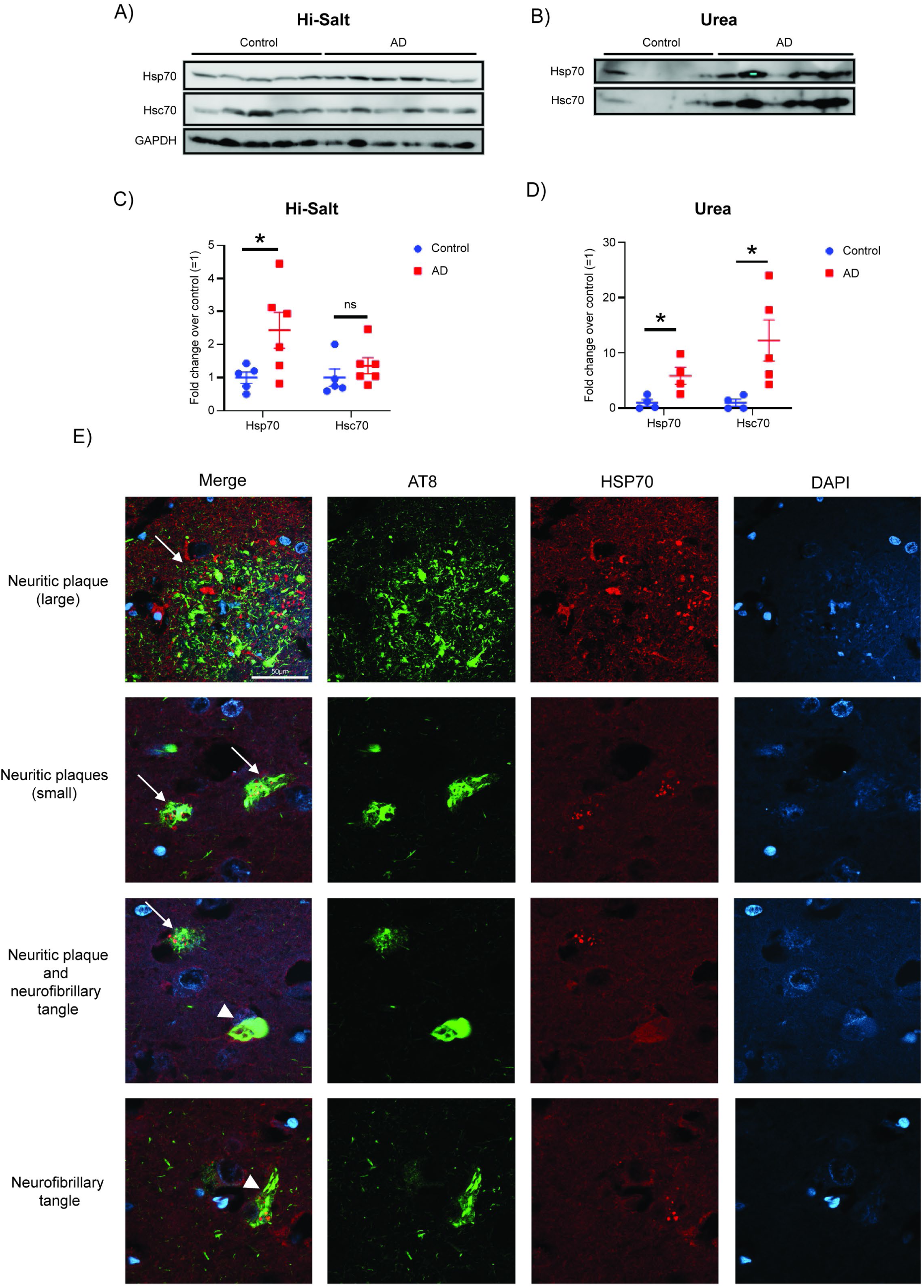
HSP70 is upregulated in human AD brain and intercalates within tau aggregates. **A)** Hi-Salt soluble immunoblots from control (n=5) and AD (n=6) cortical lysates probed for Hsp70, Hsc70, and GAPDH. **B)** Quantification of hi-salt immunoblots. *= p-value significant by unpaired Student’s t-test. **C)** Urea soluble immunoblots from control (n=4) and AD (n=5) probed for Hsp70 and Hsc70. **D)** Quantification of urea immunoblots. *= p-value significant by unpaired Student’s t-test. Overexposed Hsp70 AD sample in urea was excluded from quantification. **E)** Representative 63X confocal single-plane z-stacks from AD human hippocampus stained with AT8 (p-tau) and Hsp70. White arrow label neuritic plaques and white arrowheads label NFTs.

## Discussion

Here, we developed and characterized a cell-autonomous model of tau aggregation based on the synergistic aggregation of tau missense mutations at residues P301 and S320. Given the rapid and robust tau aggregation observed within days (in primary neurons), the PL-SF model readily lends itself to identifying factors that modulate tau aggregation. We leveraged this model to clarify the longstanding debate on the role of Hsp70 as a tau modifier. The PL-SF model could address the stepwise development of tau pathology and provide a more reliable and sensitive system for identifying subtle regulators of tau pathogenesis.

### Synergistic tau mutants as a model of mature tauopathy

Prior studies have suggested that combining tau mutations at P301 and S320 (e.g. PL-SF) can lead to accelerated tau aggregation^8–11,14^, leading to the development of new cell culture and mammalian models that better recapitulate tau pathology. For example, the S320F mutation was introduced in P301S/S305N knock-in mice, resulting in enhanced tau pathology^42^. Recently, AAV-mediated delivery of PL-SF tau to rhesus macaques was able to induce rapid and progressive tau pathology, demonstrating its utility as an efficient primate model for studying AD^43^. Here, we analyzed the biochemical and histological properties of PL-SF in cells, cortical neurons, and organotypic brain slice cultures. Strikingly, PL-SF tau inclusions appear quite similar to those detected in post-mortem AD brain (**Figures 1, 2, 6, 7**). Future cryo-EM studies may be needed to better define their ultrastructural properties^44–47^. However, if PL-SF does indeed form AD-like filaments, it may be a preferred model over other single tau variants (P301S and P301L) that lack mature amyloid structure in culture and mouse models^48,49^. The PL-SF tau variant also accumulated at the post-synaptic density (PSD), leading to functional deficits in neuronal firing and connectivity (**Figure 5 A-H**), which supports its synaptic toxicity. This rapidly maturing pathology likely allows for sensitized functional defects at the synapse.

Exactly how these two mutations (P301L or S320F) synergize to promote tau pathology and drive tau-mediated toxicity is unclear. But we can envision several scenarios. First, P301L and P301S have been well-characterized for disrupting MT dynamics and promoting tau aggregation; however, less is known about S320F, which is linked to Pick’s disease. Recently, S320F was found to stabilize a hydrophobic pocket of tau, exposing the ^306^VQIVYK^311^ amyloid motif and thereby promoting aggregation^5^. The combination of these two mutations likely amplifies their proclivity towards β-sheet self-assembly. To further test this, we generated PL-SF I277P/I308P containing two amyloid-breaking proline mutations within the MTBR. Indeed, insertion of β-breaking mutations suppressed tau aggregation in primary neurons (**Supplemental Figure 4**), in line with previous in vitro studies^37–41^.

Second, by extension of its aggregation propensity, PL-SF tau may also be more highly decorated with pathological PTMs than either single tau mutant, driving conformational changes (i.e., MC1). In this study, we only evaluated well-known tau phosphorylation epitopes (AT8, AT100, AT180), but future mass spectrometry approaches could determine the temporal emergence of global tau PTMs (**Figure 2A, B**). For instance, increased PTM complexity could create a feed-forward cycle leading to further loss of tau-MT binding. One can also envision PTM-induced loss of tau binding to quality control machinery, which typically acts in a compensatory manner to prevent aberrant tau^7^. One example is the surveillance factor HDAC6, which binds and maintains tau in the deacetylated state^7^. Thus, PL-SF could overburden the cellular proteostasis capacity, as suggested for human AD^50–52^.

Third, P301L and S320F mutations are also known to dissociate tau from microtubules, which would shift and favor their equilibrium, further concentrating tau pools in the cytosol and facilitating their aggregation. PL-SF drastically reduced acetylated-tubulin levels back down to baseline (i.e., non-tau expressing cells) (**Figure 1 H, I**), suggesting PL-SF is almost completely disassociated from microtubules and therefore more readily primed for aggregation.

Lastly, once formed, we found that neuronal PL-SF tau can recruit endogenous mouse tau (detected by T49 antibody) into tau inclusions (**Figure 2H**), consistent with a prior study showing that human PL-SF-expressing mice (SPAM mice) recruited endogenous mouse tau^11,14^. PL-SF tau, therefore, may be more highly seeding competent, in a species-independent manner, which could be leveraged to model tau spreading since it poses several advantages over the use of exogenously added, brain-derived AD seeds (e.g., large-scale production, reliability and reproducibility, and fully intact N-terminal domains). We reiterate that further ultrastructural studies are needed to determine whether seeding-competent PL-SF resembles human AD brain-derived tau.

### The role of Hsp70 in tau pathology

We added clarity to the long-standing question of whether Hsp70 modulates tau aggregation, as the prior literature is inconsistent and often highlights contradictory models for tau triage via coordinated Hsp binding. Some studies have found that activation or overexpression of Hsp70 can suppress tau aggregation in vitro, which would intuitively suggest a classical Hsp70 chaperone activity as a suppressor of tau aggregation^17,32,33,53^. Adding additional complexity, while Hsp70 can disassemble tau filaments isolated from human AD brain, this chaperone activity can release seeding-competent tau species, suggesting any protection afforded by Hsp70 may depend on the nature of the chaperone-liberated tau species^54^. In contrast, other studies have shown that Hsp70 depletion or small molecule inhibition (YM-01, JG-23/48) may drive tau clearance^22,23,28,30,55–57^. We sought to reconcile these differences in the context of our PL-SF cellular model of tau aggregation.

We initially found that Hsp70 was tightly associated with PL-SF tau aggregates in vitro and in cell culture, a finding consistent with a previous study^58^ (**Figure 5I**). HSP70 overexpression was associated with PL-SF phosphorylation and aggregation, while HSP70 depletion reduced total tau levels, including hyperphosphorylated and aggregated tau species (**Figure 3A-D**, **Figure 4A-D**). Furthermore, abrogating Hsp70 binding using an Hsp70-binding deficient mutant (PL-SF-4Δ) was also sufficient to suppress tau aggregation (**Figure 1, 2, 5, 6)**. Further supporting a role for Hsp70 in human AD pathology, we found that Hsp70 is upregulated in AD hippocampus and heavily intercalated within neuritic plaques and NFTs (**Figure 7**). The Hsp70 localization pattern was highly consistent in human AD hippocampus (**Supplemental Figure 8**) and illustrates a unique tau-Hsp70 pattern of co-pathology^59–61^. The lack of complete colocalization suggests that Hsp70 recruitment to tau occurs in a time-dependent manner. At least in a mature tau aggregation model, these data are consistent with Hsp70 stabilizing a misfolded tau conformation that potentially evades proper triage and/or degradation.

In addition to Hsp70, tau is also an Hsp90 substrate, which competes for binding sites within tau’s MTBR and is thought to promote CHIP-mediated tau degradation and clearance^21,25,62–67^. It is plausible that aggregated tau may form as a consequence of an imbalance in the ratio of Hsp70 vs. Hsp90 binding, which, if shifted towards Hsp70, could radically shift tau processing away from degradation towards stabilization of aggregated tau intermediates. In this scenario, Hsp70 depletion or inhibition would facilitate proteasome or autophagic targeting (degradation) by shifting the balance away from tau refolding. Future proteomics studies will be needed to determine whether distinct tau triage complexes emerge temporally in response to tau aggregation. For example, identifying the PL-SF interactome over time may reveal distinct triage stages in which Hsp70 has shifted from pro-folding to pro-aggregation^19^.

We did not observe any effect of Hsc70 on tau aggregation. Though highly homologous, Hsc70 is constitutively expressed at ∼30-fold higher levels in the brain compared to the inducible Hsp70 isoform^16^. Hsp70, but not Hsc70, overexpression increased the number of MC1-positive neurons by nearly 3-fold (**Figure 3E, F**). Since Hsp70 is more sensitive to redox signaling (via cysteine residue reactivity), its activity towards tau may be more tunable under stress conditions^68^. Prior work has shown that the functional outcome of Hsp70-tau interactions is highly dependent on the specificity and fine-tuning provided by co-chaperones that regulate Hsc70^34^, Hsp90^69–71^, CHIP^16^, as well as small heat shock proteins^72–74^. Thus, we suspect that many variables could dictate how effectively and efficiently Hsp70 targets tau, including: 1) the extent of MT-dissociated aggregated tau present, 2) the relative balance and abundance of chaperones, co-chaperones, and cofactors that target and triage tau, and 3) the notion that disease onset and disease progression (either acute or chronic) may involve distinct signaling pathways or compensatory responses.

Taken together, our study supports the notion that depletion or activity-modifying approaches targeting Hsp70 (or its cofactors and co-chaperones) may be neuroprotective in the context of the AD brain. While brain-specific HSP70 knockout mice are not available (to the best of our knowledge), brain-permeable Hsp70 inhibitors may have therapeutic utility, although broad Hsp70 inhibition could have detrimental effects on overall brain proteostasis. Therapeutic strategies aimed at “precision proteostasis” could selectively modulate chaperone–client interactions to favor the clearance of pathogenic tau without compromising global protein homeostasis and without directly targeting tau itself.

## Materials and Methods

### Plasmid and lentivirus cloning

To generate the pUltra-T40 WT, PL, PL-SF, SF, PL/SF-2Δ (277-278), PL/SF-2Δ (308-309), PL/SF-4Δ, HSC70-flag, HSP70-flag, and HSP90-flag constructs, the cDNAs were amplified through PCR and inserted into the pUltra vector using AgeI and SalI restriction endonucleases to replace eGFP in pUltra. pUltra-XeGFP was made by removing eGFP using AgeI and BspEI, creating a blunt end with Klenow before circularizing. pUltra-T40 WT (eGFP), PL/SF (eGFP), and PL/SF-4Δ (eGFP) were created by replacing TDP43 in pUltra TDP43 WT (eGFP) using BspEI and SalI. To generate PL/SF/2IP, two consecutive Q5 site-directed mutagenesis were performed on PL/SF(eGFP). pUltra was ordered from Addgene (Plasmid #24129). pUltra was a gift from Malcolm Moore (Addgene plasmid # 24129; RRID:Addgene_24129). shRNA plasmids against HSP70 (TRCN0000098597, TRCN0000098598) and HSP90 (TRCN0000008491, TRCN0000008494) were ordered from the UNC Lenti-shRNA Core Facility. Primers used for vector construction are in Table 1.

**Table 1:**
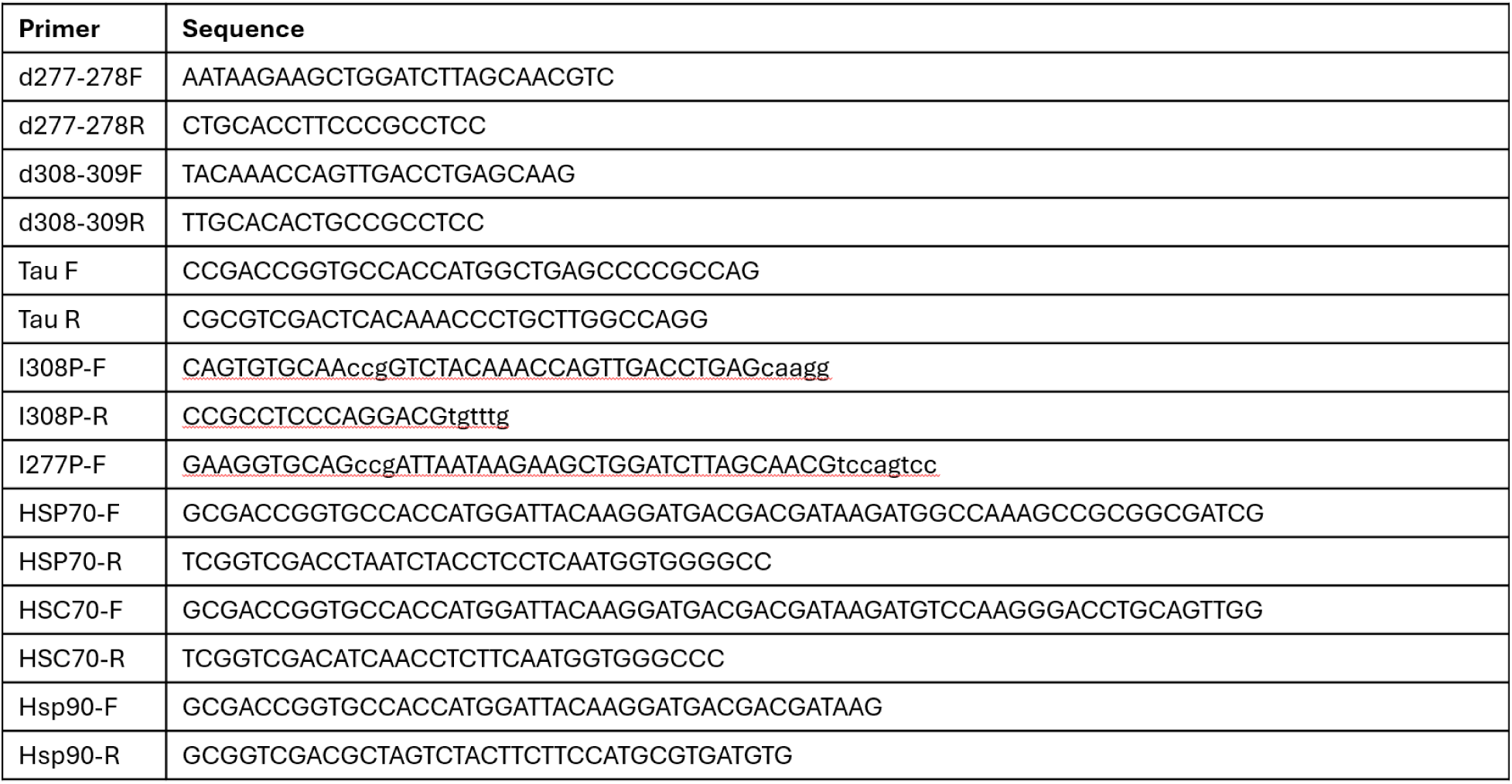
Primers used for vector construction.

### QBI-293 cell culture

QBI-293 (HEK293A) cells (ThermoFisher, #R70507) are commercially available and maintained according to standard protocols. QBI-293 cells were cultured in complete DMEM media (10% fetal bovine serum, 1X L-glutamine, and 1X penicillin/streptomycin). QBI-293 cells were transfected with FuGENE 6 transfection reagent (Promega) to express 2N4R tau constructs for 48 hr. Following transfection, cells were lysed and collected in 1X RIPA buffer [50 mM Tris pH 8.0, 150 mM NaCl, 5 mM EDTA, 0.5% sodium deoxycholate, 1% Igepal CA-630, 0.1% SDS] supplemented with deacetylase, phosphatase and protease inhibitors: 1 mM phenylmethylsulfonyl fluoride (PMSF), 10 mM Nicotinamide (NCA), 1 mM sodium orthovanadate (Na3VO4), 1 mM sodium fluoride (NaF), 0.5 M β-glycerophosphate, 2 μM Trichostatin A (TSA), and protease inhibitor cocktail composed of L-1-tosylamide-2-phenylethylchloromethyl, N-tosyl-L-lysine chloromethyl ketone, leupeptin, pepstatin, and soybean trypsin inhibitor, each at 1 μg/ml.

### Primary cortical neuron cell culture

Mouse primary cortical culture was performed in accordance with the University of North Carolina (UNC) Institutional Animal Care and Use Committee (UNC IACUC protocol 24-190). Mouse cortical neurons were dissected and isolated from C57Bl/6 mice (Charles River) at embryonic day 15-16 following lethal isoflurane anesthesia. Following anesthesia, embryos were removed from the pregnant abdominal cavity, washed well with ice-cold 70% ethanol and placed in cold Hanks-buffered balanced salt solution (HBSS). Once rinsed, these embryos were moved to a 10cm dish for dissection. The embryos were then removed from the placental membrane and brains were isolated. The cortex was dissected then moved to Hibernate-E solution (BrainBits) with B27 (Gibco) and Glutamax (ThermoFisher). The cerebral cortices were minced thoroughly with forceps and incubated for 30 minutes at 37 **°**C in HBSS with 20 U/mL papain (Worthington Biomedical), 1mM EDTA, 0.2 mg/mL L-Cysteine, and 5U/mL DNAse (Promega). Cortical tissue was further dissociated in plating media (BrainPhys media (Stemcell), 5% fetal bovine serum, 1X B27 (Gibco), 1X penicillin/streptomycin (Gibco), and 1X Glutamax by triturating the sample using a P1000 micropipette. This suspension was centrifuged for 5 minutes at 1.5 rcf to pellet the cells. The pellet was then resuspended in plating media and passed through a 40 μm cell strainer. A Countess 3 automated cell counter (Invitrogen) was used to count the cells. Cells were plated in plating media at 600,000 cells/well in 6-well plates or 30,000 cells/well in 96-well plates. The next day (DIV1), plating media was completely aspirated and replaced with fresh neuronal media (BrainPhys media (Stemcell), B27 (Gibco), 1X penicillin/streptomycin (Gibco), and 1X Glutamax).

### Lentiviral production

Lentiviral production was performed by co-transfecting 37.5 mg lentivirus plasmid with 25 mg psPAX2, 12.5 mg VSVG, and 6.25 mg REV plasmids per 15 cm culture plates of Lenti-X 293 T cells (Takara). For each lentiviral construct, three 15 cm plates were used for production. Roughly 48 hrs after the transfection, cell culture media was collected and subsequently centrifuged at 2000 rcf for 10 minutes. Lentiviral particles were purified using a double-sucrose gradient method as follows: supernatants were loaded onto a 70–60% to 30–20% sucrose gradient and centrifuged at 70,000 rcf for 2 hr at 17 °C. The 30–60% fraction containing the viral particles was retrieved, resuspended in PBS, filtered with a 0.45 mm filter flask, loaded onto a 20% sucrose solution, and centrifuged a second time at 70,000 rcf for 2 hr at 17 °C. The supernatants were carefully discarded, and the viral particles in the pellet were resuspended in PBS, aliquoted and stored at − 80 °C.

### Lentiviral transductions

Primary neurons were transduced with ∼3.0×10^12^ viral particles per mL at DIV4 for most experiments (viruses were added at DIV18 for MEA experiments). If neurons were co-transduced with two viruses, both were added sequentially (each at ∼3.0×10^12^ viral particles per mL). For transductions, viruses were added directly to the media in each well (1 µl virus per 2 mL media), followed by gentle rocking by hand to distribute viral particles. The following day, virus-containing media was completely aspirated and replaced with half conditioned media (neuronal media from the same neuron culture) and half fresh neuronal media (BrainPhys (StemCell) with B27 (Gibco), Pen-strep (Gibco), and Glutamax (Gibco)). Every two-three days, half of the neuronal media was discarded and replenished with fresh neuronal media until neurons were collected. Most neurons were collected at DIV10 (7 days after transduction), unless otherwise noted.

### Immunoblotting and biochemical analysis

RIPA soluble lysates (250 µl per well of a 6-well plate) collected from QBI-293 and primary cortical neuron were sonicated 20 times on ice, then centrifuged at 15,000 rcf for 20 minutes at 4 °C. The supernatant was removed from the pelleted material, leaving behind a RIPA insoluble pellet. The RIPA insoluble pellet was washed with 250 µl of RIPA lysis buffer with protease, phosphatase, and deacetylase inhibitors and centrifuged at 15,000 rcf for 20 minutes at 4 °C. This supernatant was removed and discarded and 75 µl of 1% SDS was added to each pellet then sonicated 20 times at room temperature. These samples were then centrifuged at 15,000 rcf at room temperature. The supernatant from these tubes was removed and stored in separate tubes as the SDS-soluble fraction. The RIPA and SDS-soluble supernatants were then mixed with 6X loading dye with DTT and boiled at 98 °C for 10 minutes in a PCR thermocycler (BioRad). Samples were run and resolved via SDS-PAGE on 4–20% gradient gels (BioRad). Gels were transferred to iBlot 2 Midi nitrocellulose membranes using the iBlot 2 (Invitrogen) following the standard P0, 7-minute settings. Following transfer, membranes were blocked with 2% milk in 1X TBS for 30 minutes and then incubated with primary antibody overnight at 4 °C on a rocker. The next day, membranes were incubated with HRP-conjugated anti-mouse or anti-rabbit antibodies (1:1000) for 1-2 hr at room temperature. Blots were developed with ECL Western Blotting substrate (Cytiva, Amsterdam) and imaged on a GE ImageQuant LAS4000.

### Immunocytochemistry

Immunocytochemistry experiments were performed using QBI-293 cells or cortical primary neurons plated on poly-D-lysine (PDL)-coated 96-well plates. Following either plasmid transfection (48 hr) or lentiviral transduction (7 days unless otherwise stated), cells were washed once in PBS, immediately fixed in 4% paraformaldehyde for 15 minutes, then washed once more with PBS. Following fixation, cells were then permeabilized with permeabilization buffer (0.2% Triton X-100 (Sigma) in PBS) for 20 minutes then blocking buffer (4% normal goat serum in PBS with 0.05% Tween 20) for 1 hr. Primary antibodies were added in blocking buffer and incubated overnight on a rocker at 4 °C. The following day, cells were washed 3X in PBS with 0.05% tween 20 and incubated with Alexa 488, Alexa 594, or Alexa 647 conjugated secondary antibodies at 1:1000 and DAPI for 1 hr. Cells were then washed 3X in PBS with 0.05% Tween 20. Cells were then imaged using an EVOS M7000 automated microscope with EVOS 20X objective (Invitrogen AMEP4982) and 40X objectives (Invitrogen AMEP4754) and DAPI, GFP, Texas Red, and Cy5 light cubes. Note: Some representative images were artificially brightened or contrasted in Microsoft PowerPoint or Celleste 5.0 and 6.0 software (Invitrogen) to aid visualization. Any panels assessing the same set of conditions were brightened to identical levels. Images were not manipulated prior to quantification.

### Antibodies

The following primary antibodies were used in this study: AT8 (ThermoFisher Scientific-MN1020), AT100 (ThermoFisher Scientific MN1060), AT180 (ThermoFisher Scientific MN1040), TauC3 (ThermoFisher Scientific-AHB0061), T49 mouse tau (Millipore-MABN827), Tau12 (Millipore, MAB2241, 1:1000), Tau5 (AHB0042), Hsp70 used for immunoblotting (Enzo-ADI-SPA-810-D), Hsp70 used for immunoblotting and immunocytochemistry (Cell Signaling Technologies-4872), Hsc70 1B5 (ADI-SPA-815-F), Hsp90 16F1 (Enzo-adi-spa-835), Total tau rabbit polyclonal-formerly K9JA (DAKO, A0024 1:2000), GAPDH 6C5 (Millipore-MAB374), ac-tubulin (Sigma T7451), alpha tubulin (Sigma, T9026), Anti-FLAG M2 (Sigma, F1804, 1:2000), GFAP (Dako, 20334). The MC1 antibody was kindly shared by Dr. Peter Davies. Quantification of immunoblotting was measured by ImageStudio Lite Version 5.2.

### Co-immunoprecipitation reactions

All co-immunoprecipitations were performed in QBI-293 cells transfected with plasmids expressing 2N4R wild-type, P301L, P301L/S320F, and P301L/S320F 4Δ for 48 hrs prior to collection. Each well of a 6-well plate was used as separate conditions. Cells were collected in 250 µl RIPA buffer with protease, phosphatase, and deacetylase inhibitors prior to co-immunoprecipitation. Lysates were immediately rotated on a wheel for 30 minutes at 20 rpm at 4 °C. Lysates were then centrifuged at 15,000 rcf for 15 minutes at 4 °C. Supernatants were removed and 40 µl were kept as input. The remaining supernatant was pre-cleared with 50 µl of Protein A/G PLUS-Agarose beads (Santa Cruz, sc2003) for 30 minutes at 4 °C. Lysates were centrifuged for 1 minute at 1,500 rcf, and the supernatant was removed. The volume of each reaction was brought to 500 µl of RIPA buffer with inhibitors. For each reaction, 8 µl of tau5 antibody was added to each reaction, then 75 µl of agarose protein A/G beads. Lysates were rotated on a wheel at 20 rpm at 4 °C overnight. The following day, lysates were centrifuged at 1,500 rcf for 5 minutes. Most of the supernatant was carefully removed, followed by washing with 500 µl of RIPA buffer with inhibitors, and lysates were centrifuged at 1,500 rcf for 5 minutes. This wash was repeated four times. After the final wash, loading dye with DTT was directly added to the beads and heated for 10 minutes at 98 °C with intermittent mixing. These were centrifuged for 30 seconds at 1,500 rcf prior to storage at -80 °C.

### Purification of recombinant tau

Tau purification methods were adapted from Barghorn et al, 2004^75^. Untagged wild-type and mutant versions of tau were cloned into pRK172 backbone and transformed into BL21(DE3)-RIL cells. Cells were grown in LB media at 37 °C until the OD600 reached 0.8. Protein expression was then induced with IPTG and continued shaking at 37 °C for 3-5 hrs. After the cells were harvested and resuspended in lysis buffer (20 mM MES pH 6.8, 1 mM EDTA, 5 mM DTT, protease inhibitors benzamidine and leupeptin, and PMSF), the cells were lysed via sonication. Lysed cells were heat shocked at 80 °C for 20 minutes, then centrifuged at 17,500 rpm (36,635 xg) for 30 minutes to remove cell debris. Clarified lysate was dialyzed overnight at 4 °C into wash buffer (20 mM MES pH 6.8, 50 mM NaCl, 1 mM EDTA, 1 mM MgCl_2_, 2 mM DTT). Following overnight dialysis, lysates were centrifuged at 17,500 rpm (36,635 xg) for 30 minutes to remove precipitated proteins and then subjected to cation exchange chromatography. The protein was eluted by increasing salt concentration in buffer to 250 mM. The samples were further purified via size exclusion chromatography on a Superdex 200 Increase 10/300 GL column in sizing buffer (25 mM HEPES pH 7.5, 100 mM NaCl, 5 mM MgCl_2_, 1 mM DTT). Samples were then concentrated as needed using centrifugal filters and flash frozen for later use.

### Heparin-induced tau aggregation assays

Tau aggregation assays were carried out in a PolarStar Omega microplate reader using Thioflavin T (ThT) fluorescence to monitor formation of tau aggregates. Aggregation reactions were started by combining 5 µM tau, 2 µM heparin, and 20 µM ThT in reaction buffer (1x PBS, 2 mM TCEP) and transferred to a costar 96-well half-area microplate (black polystyrene, flat bottom, non-treated). The microplate was covered with an ELISA plate sealer and placed in the microplate reader at 37 °C. Fluorescence measurements were taken every 5 minutes for 12 hrs using a 440-10 nm excitation and 485-12 nm emission filter.

### HSP70 and HSP90 shRNA knockdown

All shRNAs were obtained from UNC Lenti-shRNA core facility and cloned into the pUltra vectors. Sequences: HSP70 shRNA: GCAGGTGAACTACAAGGGCGA; eGFP control shRNA: TACAACAGCCACAACGTCTAT (Y145 to Y151 residues of eGFP). HSP70 shRNAs were added to the neuron culture four days after transduction with tau lentiviruses at a titer of ∼3.0×10^12^ viral particles per mL. Media was half exchanged for fresh neuronal media every two days. After an additional four days in culture, primary neurons lysates were collected for experiments. We noted some toxicity after more than 4 days in culture with HSP70 shRNAs. HSP90 shRNAs were applied for 7 days in culture (∼3.0×10^12^ viral particles per mL) prior to collection for experiments and were not overtly toxic.

### Multi-electrode array recordings and analysis

Following dissection, primary mouse cortical neurons were directly plated onto poly-D lysine coated 24-well BioCircuit MEA plates (Axion Biosystems). A 10 µl droplet of neurons at a concentration of 16,000,0000 neurons/mL was added directly to the center of each well. Neurons were incubated at 37 °C for 1 hr then 250 µl of plating media (BrainPhys media (Stemcell), 5% fetal bovine serum, 1X B27 (Gibco), 1X penicillin/streptomycin (Gibco), and 1X Glutamax) was carefully added to each well in a circular motion. Following another 1-hr incubation, another 250 µl of plating media was added to each well (total volume = 500 µl). The following day, ∼90% of plating media was aspirated and fresh neuronal (BrainPhys media (Stemcell), B27 (Gibco), 1X penicillin/streptomycin (Gibco), and 1X Glutamax) media was added to each well. Half of the media was exchanged for fresh neuronal media every two days. At DIV18, neurons underwent an initial, 10-minute pre-recording using the Axion Maestro Edge (Axion Biosystems) and AxIS Navigator software (Axion Biosystems). Immediately following this recording, neurons were transduced with lentiviruses at ∼3.0×10^12^ viral particles per mL (four replicate wells per condition). The following day, neuronal activity was recorded for 10 minutes, then lentivirus-containing neuronal media was completely replaced with half fresh neuronal media and half conditioned neuronal media from the same culture. Neurons were recorded nearly every day for 10 minutes and half of the neuronal media was replaced every 2-3 days after recording until DIV71 (day 53 of recording). Post-hoc analysis was performed using the NeuralMetricTool (Axion Biosystems) using default analysis options. Data was exported into Microsoft Excel, then graphed via GraphPad Prism 10 for statistical analyses and graphing. Raster plots were generated from the same data using the AxIS Metric Plotting Tool. 96 well plates were plated at the same time using the same primary neurons, underwent identical media changes, and were used for immunocytochemistry at DIV29 and DIV62 in Figure 5.

### Post-synaptic density isolation

Primary mouse cortical neurons from wild-type C57/BL6 mice were plated on poly-D-lysine coated 6-well plates at 600,000 cells/well. Neurons were treated with lentivirus at DIV4 (1 6-well plate per condition) and half-fed every two days until DIV14. At DIV14, neurons were rinsed in cold PBS and all 6-wells of each plate were collected in a total of 900 µl of homogenization buffer (320 mM sucrose, 5 mM sodium pyrophosphate, 1 mM EDTA, 10 mM HEPES pH 7.4) with protease/phosphatase inhibitor cocktail 200 nM okadaic acid, 1 mM orthovanadate, 1X protease inhibitor cocktail (Roche). Neuronal lysates were homogenized by passing the lysate 12 times using a 26g needle. For each condition, 70 µl of lysate was kept and stored as whole cell lysate. The remaining lysate was centrifuged at 1,000 g for 10 minutes at 4 °C *to* generate pelleted nuclear fraction (P1) and supernatant fraction (S1). The S1 fraction was centrifuged at 15,000 g for 20 minutes at 4 °C to generate the membrane/crude synaptosome pelleted fraction (P2) and cytosolic supernatant fraction (S2). The P2 fraction was resuspended in 500 µl of ultrapure H_2_0 with protease/phosphatase cocktail. 20 µl of 100 mM HEPES was added to the 500 µl of suspension to yield 4 mM HEPES pH 7.4. These samples were then spun on a wheel for 30 minutes at 4°C then centrifuged for 20 minutes at 4 °C at 25,000 g using a Sorvall mTX 150 ultracentrifuge (Thermo Scientific) to generate pelleted synaptic fraction (LP1) and supernatant fraction (LS2). The LP1 pellet was resuspended in 250 µl mM HEPES pH7.4 using repeated trituration with a micropipette. This was then mixed with 250 µl of 1% Triton X-100 with protease/phosphatase cocktail and spun on a wheel for 10 minutes at 4 °C. This was then centrifuged in Sorvall mTX 150 ultracentrifuge (Thermo Scientific) at 32,000 g for 20 minutes at 4 °C. This pellet was resuspended in 80 μL of 50 mM HEPES pH7.4 using micropipette trituration to yield the PSD fraction. Only total homogenate and PSD fraction were processed and analyzed by SDS-PAGE.

### Imaging quantification

All immunocytochemistry (aside from organotypic slice cultures) quantification was performed using Celleste 5.0 and 6.0 software (Invitrogen). The number of inclusions were counted using the count/size function. Puncta and inclusions were manually segmented based on intensity (Segment: Manual) and size (Region: Area) thresholds for each tau probe (e.g. AT8, MC1, ThioS). These threshold settings were then applied to all images used for that specific experiment, automatically batched processed using a segment-count-export macro generated in Celleste, then exported to Microsoft Excel for subsequent analysis.

### Hippocampal slice cultures

Sprague-Dawley rats (Charles River Laboratories, Wilmington, MA) were housed and maintained in accordance with the recommendations from the Guide for the Care and Use of Laboratory Animals from the National Institutes of Health, and in accordance with an approved protocol from the Institutional Animal Care and Use Committee of the University of North Carolina-Pembroke. Brain tissue from postnatal 12-day-old rats was rapidly prepared and hippocampi positioned on a McIlwain tissue chopper for assembling slices for interface cultures as previously described^76,77^. Three-dimensional organotypic cultures of specific brain regions are well known for studying different types of physiologies and pathogenic processes^78–80^. Transverse hippocampal slices (400-µm thickness) were immediately placed in ice-cooled-buffer solution (pH 7.2) then transferred in groups of eight to nine onto the Biopore membrane of culture inserts (Millipore, Billerica, MA). The media below the culture inserts contained 50% basal medium eagle (Sigma-Aldrich, St. Louis, MO), 25% Earle’s balanced salts (Sigma-Aldrich), 25% horse serum (Gemini Bio-Products, Sacramento, CA), and defined supplements as previously described^81^. The media were initially changed 24 hrs after the slicing procedure, followed by subsequent changes every 3-4 days. The cultivated hippocampal slices were maintained under conditions of a 5% CO2-enriched atmosphere at a constant temperature of 37 °C for a duration of nine weeks.

### Lentiviral infection of organotypic cultures

Lentiviruses expressing GFP or GFP-tagged tau variants (wild-type, PL-SF, and PL-SF 4Δ tau). For each condition, 1.5 µl of lentivirus were applied onto the surface of individual organotypic slices (∼7.5×10^12^ viral particles per slice) approximately 3 hrs after the placement of the slices on Millipore inserts. Following a nine-week incubation period in vitro, both the infected organotypic slices and their corresponding controls were fixed with 4% PFA for 3-4 hrs. Subsequently, the slices were rinsed multiple times with PBS and stored in PBS at 4 °C.

### Immunofluorescence staining and imaging of organotypic brain slice cultures

The fixed organotypic hippocampal explants underwent permeabilization in PBS containing 0.5% Triton-X100 for 30 minutes. Following this, the slices were blocked with a solution of 5% BSA in PBS containing 0.1% Triton-X100 for 90 minutes at room temperature. Subsequently, the organotypic slices were incubated for 48 hrs at 4 °C with primary antibodies targeting AT8 (1/400; ThermoFisher Scientific-MN1020) or MC1 (1/400; Dr. Peter Davies, diluted in a solution of 3% BSA containing 0.1% Triton-X100. Next, brain explants were washed 3-4 times with PBS containing 0.1% Triton-X100 and then exposed to Goat anti-mouse IgG Alexa-FluorTM 568 (ThermoFisher, Waltham, MA) for 120 minutes at room temperature. Following this incubation, the organotypic slices were rinsed three times with PBS containing 0.1% Triton-X100. During the second wash, NucBlue stain (Invitrogen, MA, USA) was introduced to the PBS to promote DAPI nuclear staining. Subsequently, the slices were rinsed in PBS and were mounted onto glass slides, covered by 1.5 mm coverslips, using ProlongTM diamond antifade mountant (Thermo Fisher Scientific, NC, USA). Immunofluorescence images were captured with Andor Dragonfly spinning disk confocal using the Zyla Plus 4.2MP sCMOS (Oxford Instruments, Abingdon, United Kingdom). Image our blinded processed adopting the equal adjust using the Imaris Multidimensional visualization and analysis software (Schlieren-Zurich, Switzerland).

### Human brain fractionation

Human control and Alzheimer’s disease (AD) cortical brain tissues (Braak stages V–VI) were obtained from the Center for Neurodegenerative Disease Research brain bank at the University of Pennsylvania. Gray matter from the frontal cortex of each subject was isolated and homogenized in 3 volumes per gram (vol/g) of cold high-salt RAB buffer (0.75 M NaCl, 100 mM Tris, 1 mM EGTA, 0.5 mM MgSO₄, 0.02 M NaF, 2 mM DTT, pH 7.4), supplemented with deacetylase, phosphatase, and protease inhibitors as previously described. Homogenates were incubated at 4 °C for 20 minutes to facilitate microtubule depolymerization, followed by ultracentrifugation at 100,000 × g for 30 minutes at 4 °C. The resulting supernatant was designated the high-salt fraction. The pellet was then re-homogenized and subjected to a second round of centrifugation under identical buffer and conditions. After removal of the supernatant, the remaining pellets were homogenized in 5 vol/g of cold RIPA buffer (50 mM Tris, pH 8.0, 150 mM NaCl, 1% NP-40, 5 mM EDTA, 0.5% sodium deoxycholate, 0.1% SDS), and centrifuged once more at 100,000 × g for 30 minutes at 4 °C. Subsequent to myelin flotation, the pellets were re-extracted in RIPA buffer containing 20% sucrose to eliminate residual myelin and labeled as the RIPA fraction. Finally, the insoluble material was extracted using 1 vol/g of urea extraction buffer (7 M urea, 2 M thiourea, 4% CHAPS, 30 mM Tris, pH 8.5). Both the high-salt and urea-soluble fractions were subjected to SDS-PAGE followed by immunoblotting with antibodies targeting Hsp70, Hsc70, and GAPDH.

### Human brain immunohistochemistry and confocal microscopy

Human hippocampal tissue sections mounted on coverslips were obtained from the UNC Translational Pathology Laboratory (TPL), which oversees a cataloged and archived repository of Alzheimer’s disease (AD) brain specimens, including neuropathologically confirmed cases from UNC-affiliated donors. The 10 um sections were de-paraffinized with two 45-minute washes in xylene followed by one-minute washes in 100% ethanol (x2), 95% ethanol, 80% ethanol, 75% ethanol, water, and finally allowed to soak O/N in 1x PBS. Antigen retrieval is performed using a citrate-based antigen unmasking solution (Vector Labs, H-3300-250) in boiling Mill-Q water for 10-minutes. The slides were then washed 3×5-min with TBS, followed by 1x wash in TBS + 0.1% TritonX-100 for 20 minutes, 3×5-min with TBS, and 1×5 min in 3% Goat Serum in TBS (Southern Biotech, 0060-01). Sections were then stained 2x O/N in a humidified chamber at room temperature using anti-AT8 (1:500, Invitrogen, MN1020) and anti-Hsp70 (1:500, Abcam, ab181606). Slides were then washed 3×5-min with TBS followed by secondary staining with goat anti-mouse IgG 488 (1:1000, Invitrogen, A-11001) and goat anti-rabbit IgG 594 (1:1000, Invitrogen, A-11012) for 24 hours at 4°C in a dark, humidified chamber. Slides were then washed 3×5-min with TBS, allowed to dry, and mounted with Fluoromount-G with DAPI (Invitrogen, 00-4959-52).

Confocal microscopy was performed on an LSM980 (Zeiss) inverted laser scanning confocal microscope. The LSM980 is equipped with a motorized X, Y, stage and Z-focus with high-speed Piezo insert along with four diode lasers (405, 488, 561, and 633). Images were acquired with 1x Nyquist sampling using 63x 1.4 NA Plan-Apochromat oil objective (Zeiss) and 4 channel GaAsP detectors on Zen Blue 3.6 software. Images were acquired with 1x Nyquist sampling using 20×0.8 NA Plan-Apochromat air objective (Zeiss) and 4 channel GaAsP detectors on Zen Blue 3.6 software.

### Statistical analysis

GraphPad Prism 10 software was used for all statistical analyses. For comparisons between two groups, two-tailed unpaired Student’s t-tests were used. For multiple group comparisons, one-way ANOVA followed by Dunnet’s or Sidak’s post hoc tests were applied as appropriate. Results were polled from a minimum of n=3 independent experiments and presented as mean ± standard error of the mean (SEM). P-values < 0.05 were considered statistically significant. The number of replicates (n) and details of the specific tests applied are provided in figure legends.

## Acknowledgements

The Andor Dragonfly microscope was funded with support from National Institutes of Health grant S10OD030223. The Zeiss LSM 980 microscope was funded with support from NIH grant S10 OD032388. We would also like to thank Kinsley Adams for her expertise and assistance with the hippocampal organotypic slice cultures. We would also like to thank the University of Pennsylvania (Center for Neurodegenerative Disease Research brain bank) for providing human control and AD cortex lysates. Lastly, we would like to thank the University of North Carolina Translational Pathology Laboratory for providing AD hippocampal brain sections.

## Author Contributions

MRB designed and performed most experiments. MA, JS, and BB designed and carried out ex vivo organotypic slices experiments and imaging. KP performed all immunohistochemistry and confocal imaging of human post-mortem brain. CO assisted with co-immunoprecipitation experiments and post-synaptic density preparations. CO and JHT performed human brain fractionation and immunoblots. EP performed all protein purification and carried out the Thioflavin T aggregation assays. JMM assisted with multi-electrode array recordings and analyses. XT prepared all plasmid and lentiviral constructs for all experiments. DK and HT designed and performed the QBI293 experiments comparing various P301 and S320 tau mutants. JHT, JS, NB, and BB provided additional experimental guidance and interpretation. MRB and TJC prepared the manuscript and figures. TJC directed and supervised the study. All authors consented and contributed to the final version of the manuscript.

## Figures and Figure Legends

**Supplemental Figure 1:**
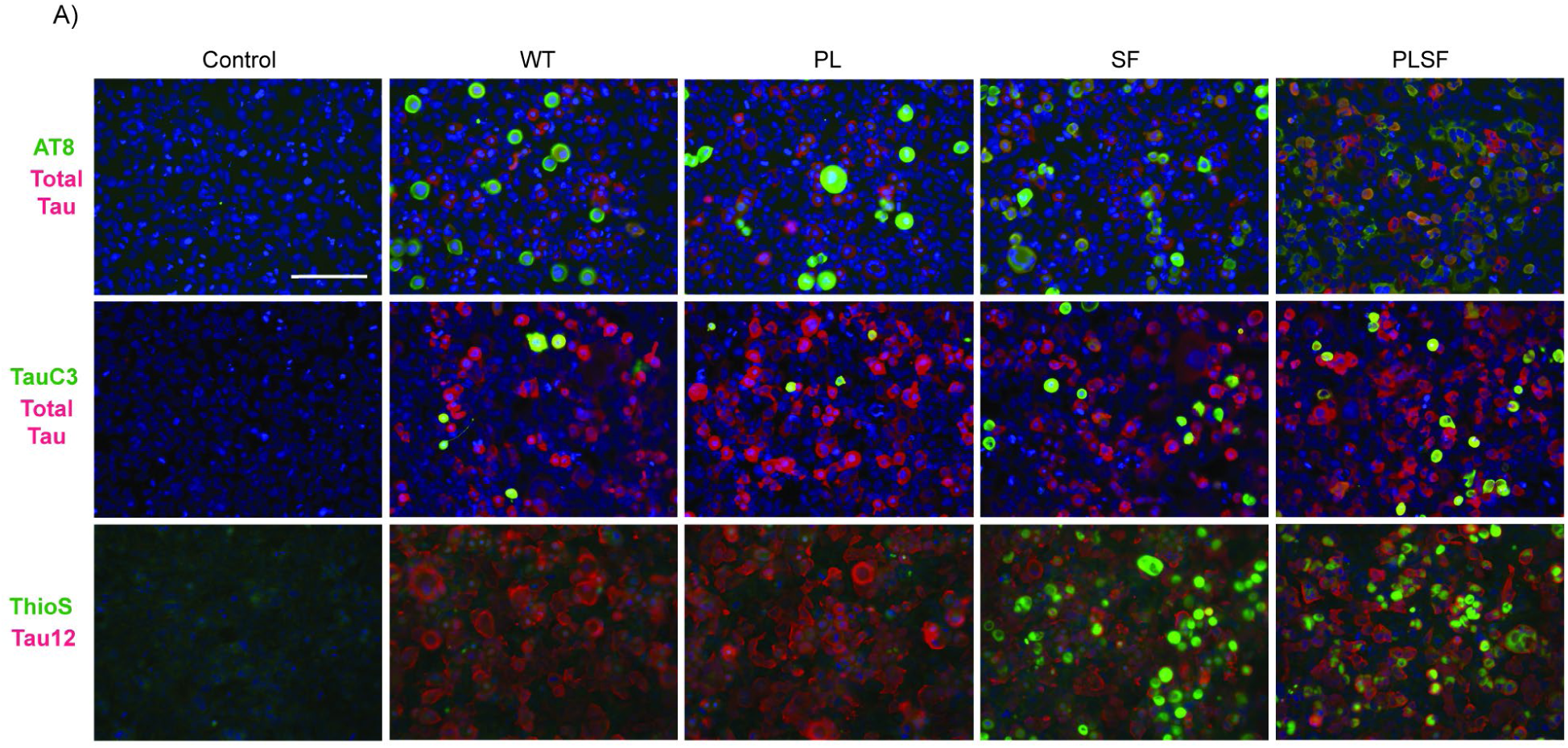
Accelerated PL-SF aggregation compared to single tau mutants in immortalized cell lines. **A)** Representative immunocytochemistry images from QBI-293 cells expressing empty vector (pcDNA) or WT, PL, SF, and PL-SF tau variants. Cells were labeled with AT8 (p-tau) or TauC3 (C-terminal cleaved tau) with rabbit polyclonal tau (total tau) and ThioS with Tau12 (human tau). Quantification of images is found in Figure 1C. Control= empty plasmid vector (pcDNA3.1). Scale bar= 125 µm.

**Supplemental Figure 2:**
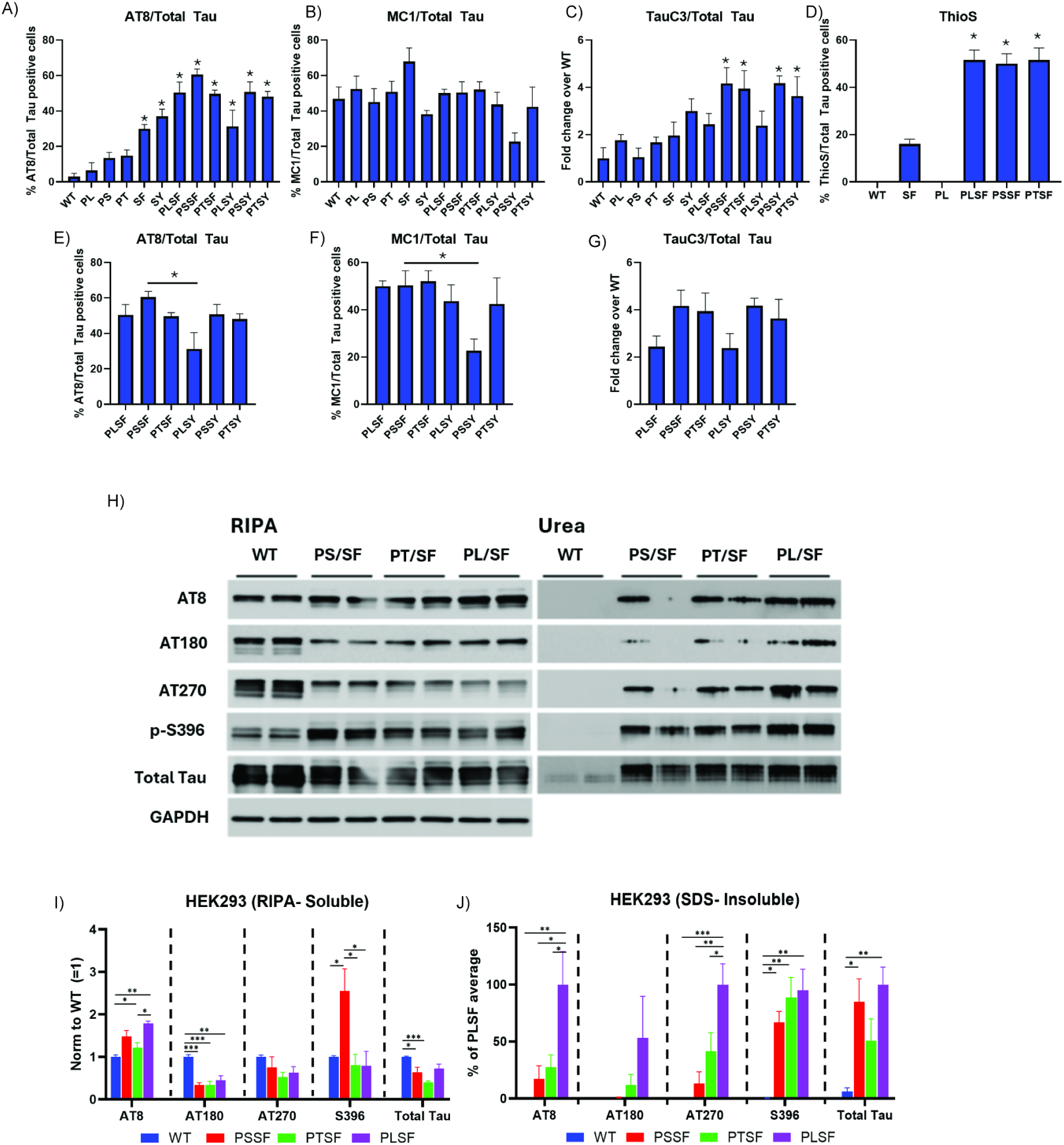
Immunoblotting analysis of a panel of P301 and S320 single and combination mutants. **A-G)** Quantification of AT8, MC1, TauC3, and ThioS immunocytochemistry in QBI-293 cells expressing various P301 and S320 mutants alone or in combination. Panels E-G are subsets of data from A-C to perform additional comparisons among combinatorial mutants. Representative images not shown. Error bars= SEM. *= significant difference by one-way ANOVA. **H)** Representative immunoblots from QBI-293 cells expressing wild-type, P301S/S320F, P301T/S320F, and P301L/S320F tau variants. RIPA (soluble) and urea (insoluble) fractions were analyzed by immunoblotting for p-tau epitopes (AT8, AT180, AT270, p-396) and total tau (rabbit polyclonal). **I, J)** Quantification of RIPA soluble and urea (insoluble) fractions. Error bars= SEM. *= significant difference by one-way ANOVA.

**Supplemental Figure 3:**
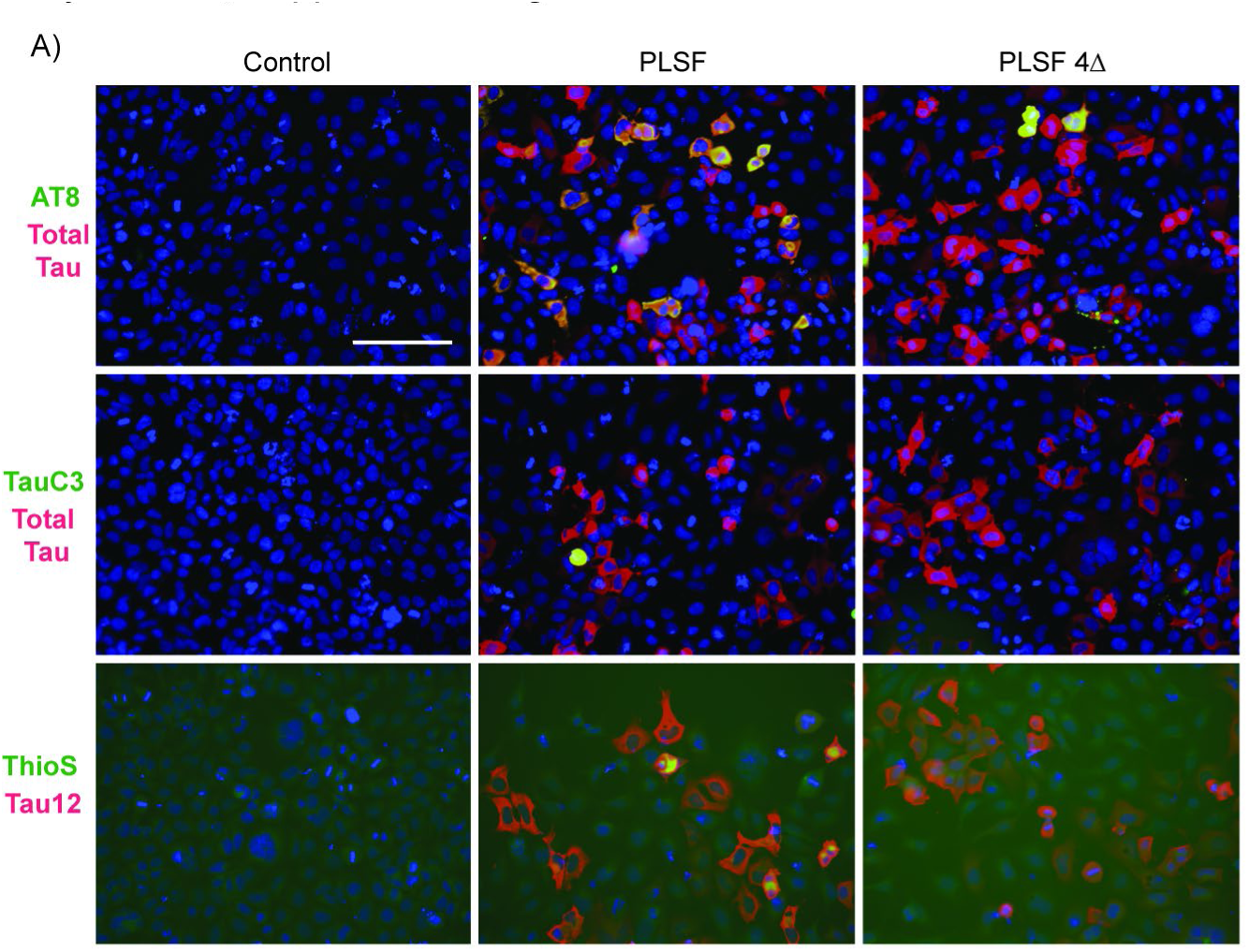
Abrogating Hsp binding via the PL-SF 4Δ variant suppresses the accumulation of pathological tau. Representative immunocytochemistry images from QBI-293 cells expressing tau PL-SF and PL-SF 4Δ tau variants. Cells were probed with AT8 (p-tau), TauC3 (C-terminal cleaved tau) with rabbit polyclonal tau (total tau), and ThioS with Tau12 (human tau). Quantification of images is found in Figure 1F. Control= empty plasmid vector (pcDNA3.1). Scale bar= 125 µm.

**Supplemental Figure 4:**
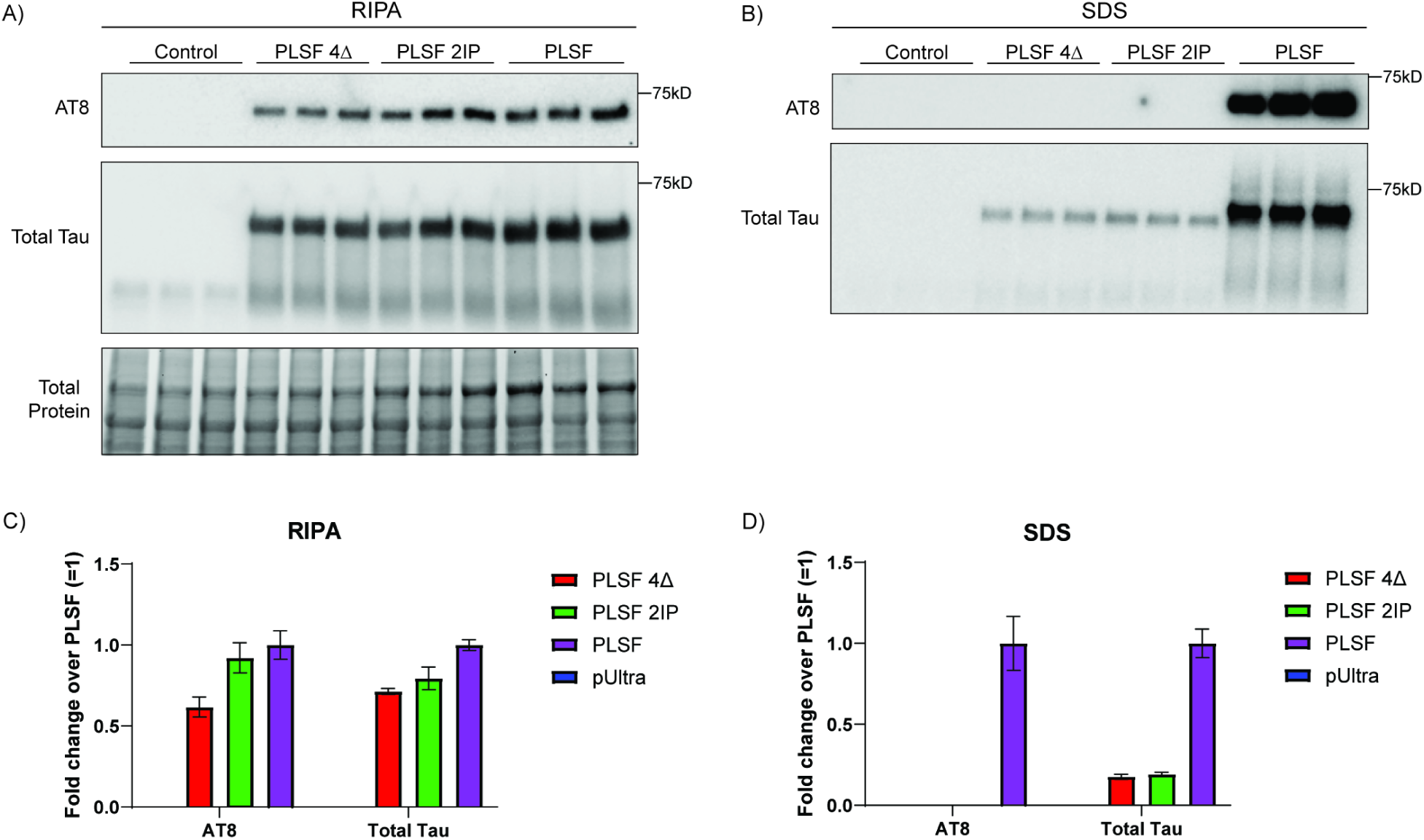
PLSF containing the β-structure breaking mutations, I277P/I308P, prevents tau aggregation similar to PLSF 4Δ. **A)** Representative RIPA immunoblot of primary neurons transduced with empty control, PLSF 4Δ-eGFP, PLSF I277P/I308P (2IP), or PLSF-eGFP and probed for p-tau (AT8) and total tau (rabbit polyclonal). **B)** Quantification of AT8 and total tau in RIPA fraction. **C)** Representative SDS immunoblot of primary neurons transduced with empty control, PLSF 4Δ-eGFP, PLSF I277P/I308P (2IP), or PLSF-eGFP and probed for p-tau (AT8) and total tau (rabbit polyclonal. **D)** Quantification of SDS fraction. *= significant difference by one-way ANOVA.

**Supplemental Figure 5:**
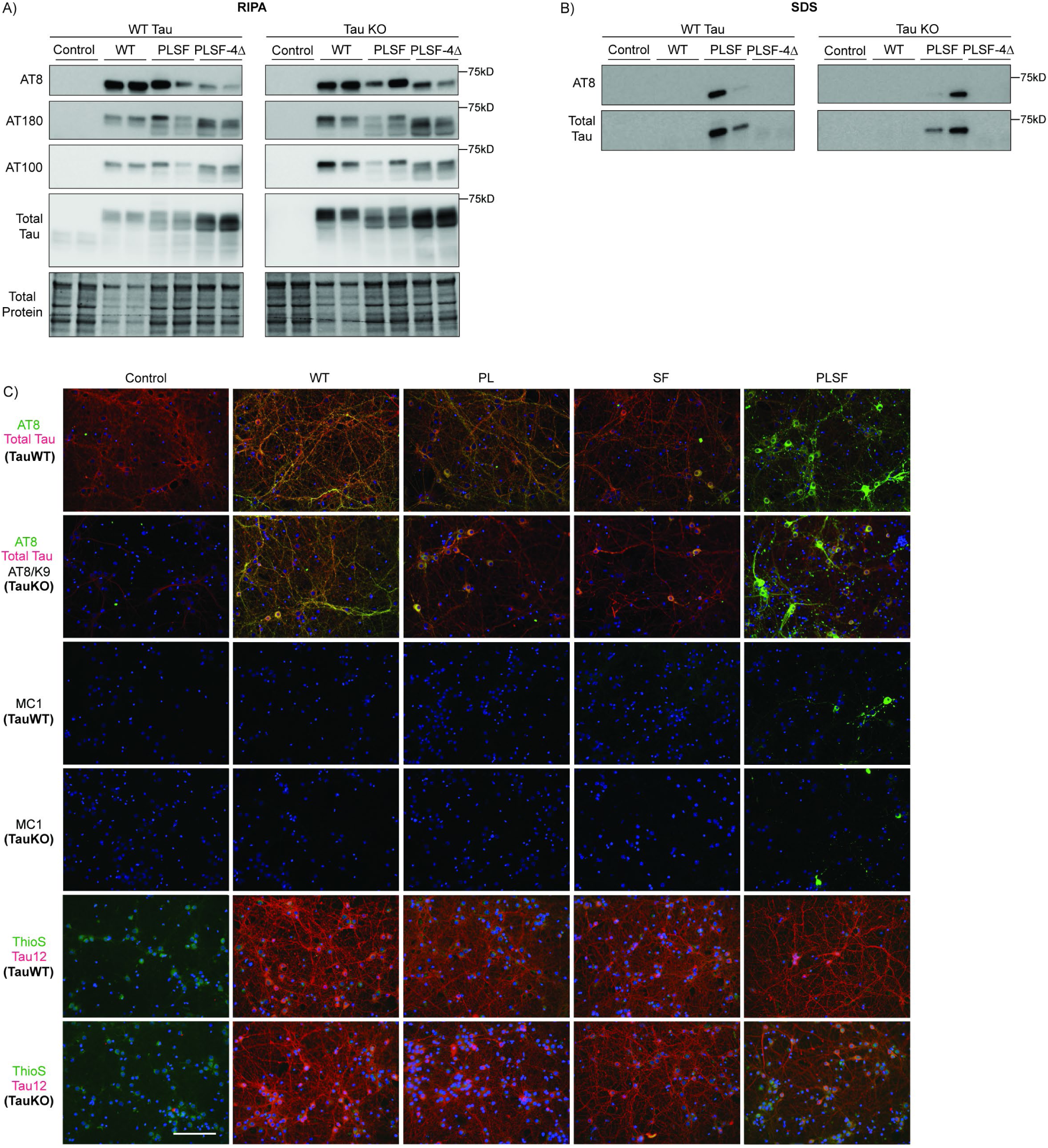
PL-SF aggregates emerge similarly in either wild-type or tau knockout neurons lacking endogenous mouse tau. **A)** RIPA (soluble) immunoblot of wild-type tau and tau knockout neurons probed with p-tau (AT8, AT180, AT100) and total tau (rabbit polyclonal). **B)** SDS (insoluble) immunoblot of wild-type tau and tau knockout neurons probed with p-tau (AT8) and total tau (rabbit polyclonal). **C)** Representative immunoblotting images of wild-type, P301L, S320F, and P301L/S320F tau variants in wild-tau-expressing or tau knockout neurons. Neurons were labeled with AT8 (GFP) and rabbit polyclonal total tau (RFP), MC1 (GFP), or ThioS (GFP) and Tau12 (RFP). Scale bar= 125 µm. Control= empty viral vector (pUltra).

**Supplemental Figure 6:**
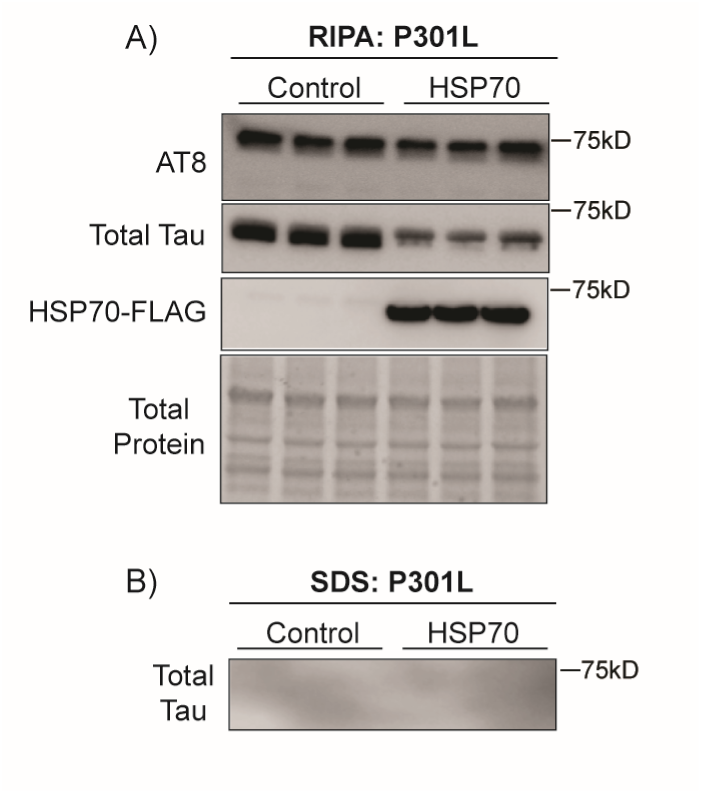
Overexpression of HSP70 in P301L-expressing neurons does not promote tau phosphorylation or aggregation. **A)** RIPA (soluble) immunoblot of primary neurons following 1-week of lentiviral co-transduction with P301L tau and HSP70. Blot was probed with p-tau (AT8), total tau (rabbit polyclonal), and M2 FLAG (Hsp70-FLAG). **B)** SDS (insoluble) immunoblot of primary neurons was unable to detect insoluble tau species with total tau antibody (rabbit polyclonal). Control= empty viral vector (pUltra).

**Supplemental Figure 7:**
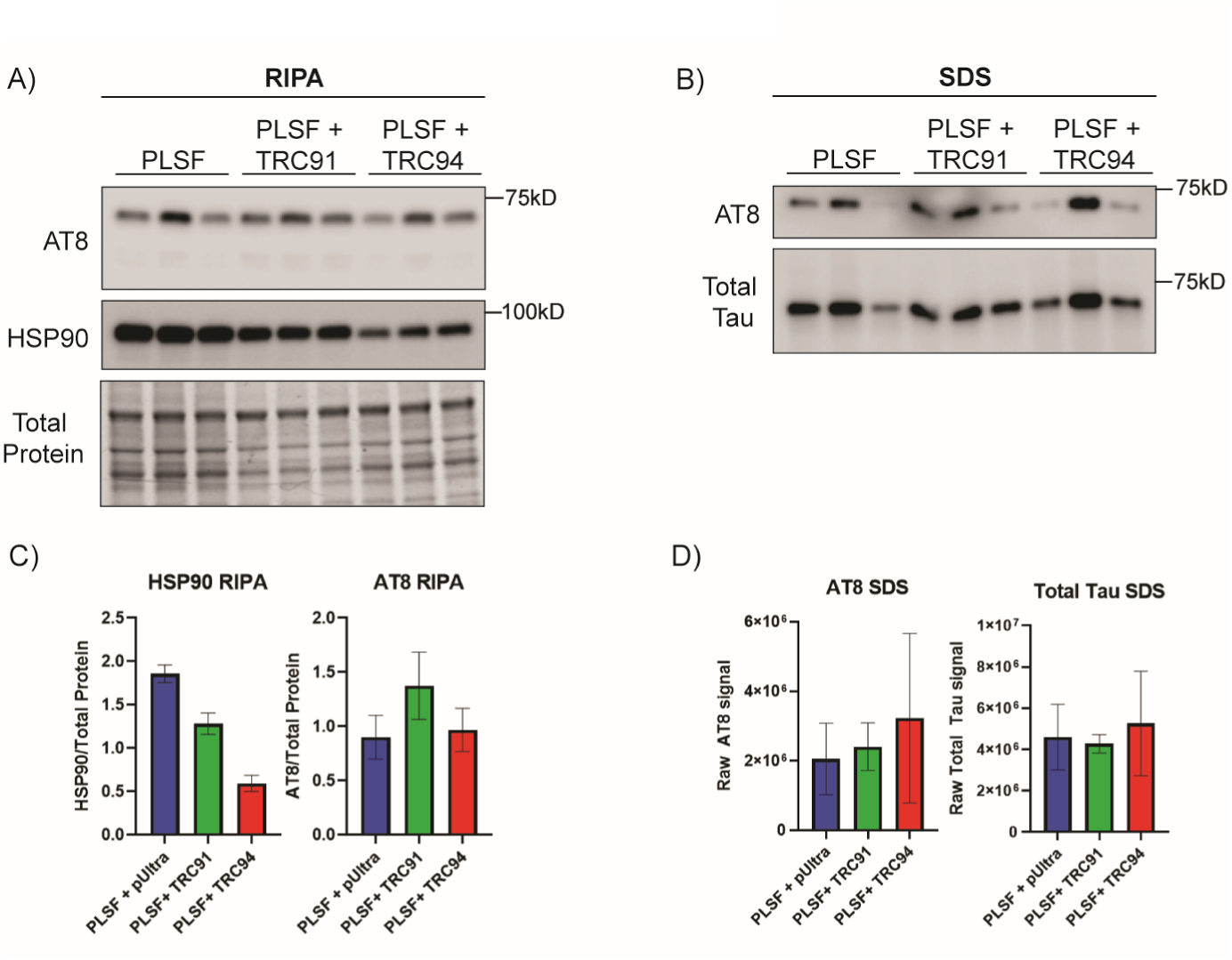
HSP90 depletion does not alter phosphorylated or aggregated PL-SF tau. **A)** RIPA (soluble) immunoblot of primary neurons probed with p-tau (AT8) and Hsp90 following 1-week of lentiviral HSP90 knockdown with two shRNAs (called TRC91 and TRC94). **B)** SDS (insoluble) immunoblot of primary neurons probed with p-tau (AT8) and total tau (rabbit polyclonal) following 1-week of lentiviral HSP90 knockdown. **C)** Quantification of AT8 and Hsp90 in the RIPA fraction. Error bars= SEM. **D)** Quantification of AT8 and total tau (rabbit polyclonal) in SDS fraction. *Note: Only TRC94, and not TRC91, HSP90 shRNA was able to reduce Hsp90 expression.

**Supplemental Figure 8:**
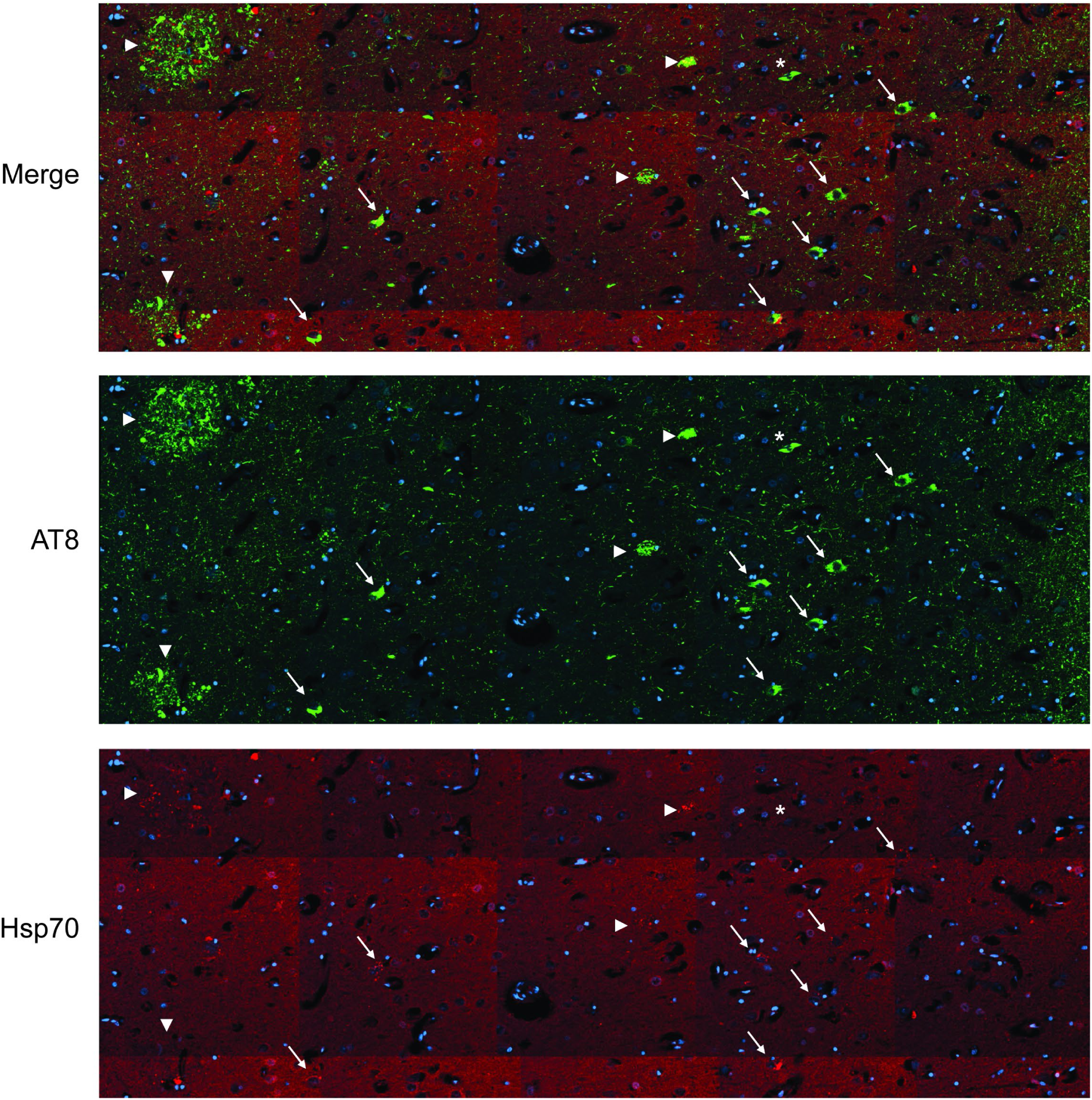
Hsp70 foci accumulation within neuritic plaques and neurofibrillary tangles in human AD brain. **A)** Representative stitched widefield images of a human AD patient hippocampus labeled with AT8 (GFP) and Hsp70 (RFP). AT8/Hsp70 positive plaques are labeled with white arrowheads. AT8/Hsp70 positive tangles are labeled with white arrows. * = a single AT8 positive tangle that is absent for Hsp70 foci.

